# A rapid, field-deployable paper-based biosensor for the detection of African swine fever virus in whole blood

**DOI:** 10.64898/2026.06.16.732725

**Authors:** Bibek Raut, Gopal Palla, Nafisa Rafiq, Jiangshan Wang, Virendra Kumar, Mohamed S. Kamel, Dan Van Nguyen, Saraswathi Lanka, Carol W. Maddox, Darryl Ragland, J. Alex Pasternak, Mohit S. Verma

## Abstract

African swine fever virus (ASFV) poses a major transboundary threat to global swine production, underscoring the need for rapid and field-deployable diagnostic tools. Although quantitative polymerase chain reaction (qPCR)-based assays are the standard molecular assay for ASFV detection, their reliance on centralized laboratory infrastructure, multi-step sample preparation, and trained personnel limit their utility for timely decision-making at the point of need (PON). Here, we report a portable molecular diagnostic platform that enables colorimetric quantitative loop-mediated isothermal amplification (qLAMP) directly from diluted whole blood on microfluidic paper-based analytical devices (µPADs). The assay targets the conserved ASFV viral protein 72 (VP72) and topoisomerase II (TOPII) genes and incorporates objective image-based colorimetric signal analysis to reduce user-dependent interpretation. Using plasmid DNA spiked into whole blood diluted to 5% (v/v) in 5% D-mannitol, the µPAD-LAMP assay achieved a limit of detection (LOD) of 25 copies per reaction (67 copies/µL of whole blood sample) for VP72 targets with no observed cross-reactivity against nine common swine pathogens, demonstrating 100% analytical sensitivity and specificity during in-house testing and 90% and 92% analytical sensitivity and specificity respectively in an external laboratory evaluation. The complete assay was performed within 60 minutes using a portable heating and imaging platform. Together, these results demonstrate a simple, DNA extraction-free molecular diagnostic approach that enables rapid and reliable ASFV detection from whole blood applicable to field-relevant conditions.

## Introduction

Although the World Organization for Animal Health (WOAH) recommends quantitative polymerase chain reaction (qPCR) as the reference molecular test for African swine fever virus (ASFV) detection, its reliance on multi-step whole-blood preprocessing limits field deployment (WOAH, 2022, 2024; King et al., 2003; Fernández-Pinero et al., 2013; Hu et al., 2023). To address this limitation, we developed a paper-based, colorimetric quantitative loop-mediated isothermal amplification (qLAMP) biosensor that delivers diagnostic results within 60 min, directly from minimally processed, extraction-free swine whole-blood samples.

ASFV poses a major transboundary threat to swine production, with the 2018–2019 outbreaks in China alone estimated to have caused more than USD 100 billion in total economic losses (You et al., 2021). ASFV is a large double-stranded DNA virus of the *Asfarviridae* family that is highly contagious in both domestic and feral pigs and can cause mortality rates up to 100% in peracute and acute cases (Tulman et al., 2009; Dixon et al., 2020; Le et al., 2024; Solikhah et al., 2025). Although ASF was historically confined to sub-Saharan Africa, it has spread globally over the past two decades, with major incursions into Eastern Europe in 2007, East Asia including China in 2018, and the Caribbean in 2021 (Eustace Montgomery, 1921; Rowlands et al., 2008; Zhou et al., 2018; Dixon et al., 2020; Schambow et al., 2025). While ASF has not been detected in the United States, the world’s third-largest pork producer, the close geographic proximity of ongoing Caribbean outbreaks and the absence of a vaccine licensed for use in the United States underscore the ongoing risk to the U.S. pork industry and the need for effective preparedness and early-detection strategies (Brown et al., 2024; Lim et al., 2023; See, 2024; Sykes et al., 2025).

Early diagnosis through laboratory testing is critical for effective control of ASF because acute infection progresses rapidly and clinical signs cannot be reliably distinguished from other febrile hemorrhagic syndromes or bacterial septicemias of pigs by clinical or post-mortem examination (WOAH, 2022; Lim et al., 2023; Hu et al., 2023; Ekakoro et al., 2025). Rapid detection is also essential for biosecurity and border surveillance, as ASFV can persist in contaminated pork and pork products, which are recognized pathways for long-distance transboundary spread through trade and illegal importation of meat products (Arcega Castillo et al., 2025; Ratnawati et al., 2025; Thakur et al., 2026). Accordingly, the WOAH recommends qPCR as the standard molecular assay for ASFV detection and confirmation (WOAH, 2024). In particular, the internationally harmonized King and Universal Probe Library (UPL) assays targeting the highly conserved viral protein 72 (VP72) capsid protein (also called p72) region are widely used for detection of acute infections (King et al., 2003; Fernández-Pinero et al., 2013; WOAH, 2024). In addition to these assays, WOAH recognizes several other validated qPCR assays, and numerous studies report molecular tests with analytical sensitivity and specificity approaching 100% and no cross-reactivity with other porcine viruses and bacteria (Gu et al., 2025; Hu et al., 2023; Zhang et al., 2025; Zhu et al., 2024). Despite the availability of these highly accurate laboratory diagnostics, standard qPCR workflows rely on centralized laboratory infrastructure, sample submission to reference laboratories, and multi-step sample preparation steps (e.g., dilution, DNA extraction and purification) performed by trained personnel, which delays actionable results and constrains rapid decision-making at farms, markets, and outbreak checkpoints (Arcega Castillo et al., 2025; Ratnawati et al., 2025; Thakur et al., 2026; WOAH, 2022).

To address the operational constraints of laboratory-based qPCR workflows, substantial effort has focused on adapting molecular assays for use closer to the point of need (PON) in application ranging from animal health (Davidson et al., 2026; Kamel et al., 2025a, 2025b; Mohan et al., 2021; Pascual-Garrigos et al., 2021), environment monitoring (Ranjbaran et al., 2024; Wang et al., 2023), and human health (J. Wang et al., 2021a, 2021b). In particular, isothermal amplification methods, most notably LAMP (Notomi et al., 2000; Mori and Notomi, 2009), have attracted considerable interest because they operate at a constant temperature (60-65 °C) and can support simplified workflows with fluorescence or visual readouts using minimal instrumentation for heating and optical detection (Tran et al., 2021; Bohorquez et al., 2023; Zhang et al., 2025; Gu et al., 2025; Davidson et al., 2026; Kamel et al., 2025b; Rafiq and Verma, 2024; Raut et al., 2026a). In addition to LAMP, a wide range of PON systems based on portable PCR, alternative isothermal amplification methods (e.g., recombinase polymerase amplification (RPA) and recombinase-aided amplification (RAA)), and more recently clustered regularly interspaced short palindromic repeats (CRISPR)-based nucleic acid detection have been reported and summarized in several recent review articles (Gu et al., 2025; Hu et al., 2023; Lim et al., 2023; WOAH, 2022; Zhang et al., 2025; Zhu et al., 2024). From these studies, we selected the most relevant LAMP-based PON systems published within the past five years (since 2020) and summarized them in Table 1 for comparison with our system. Across these reported LAMP assays, commonly targeted genomic regions include VP72 (also called p72, gene *B646L*) and topoisomerase II (TOPII), with demonstrated limits of detection (LOD) down to 10 copies/µL in serum sample spiked with plasmid DNA (Tran et al., 2021) and time-to-result of less than 60 min.

**Table 1.**
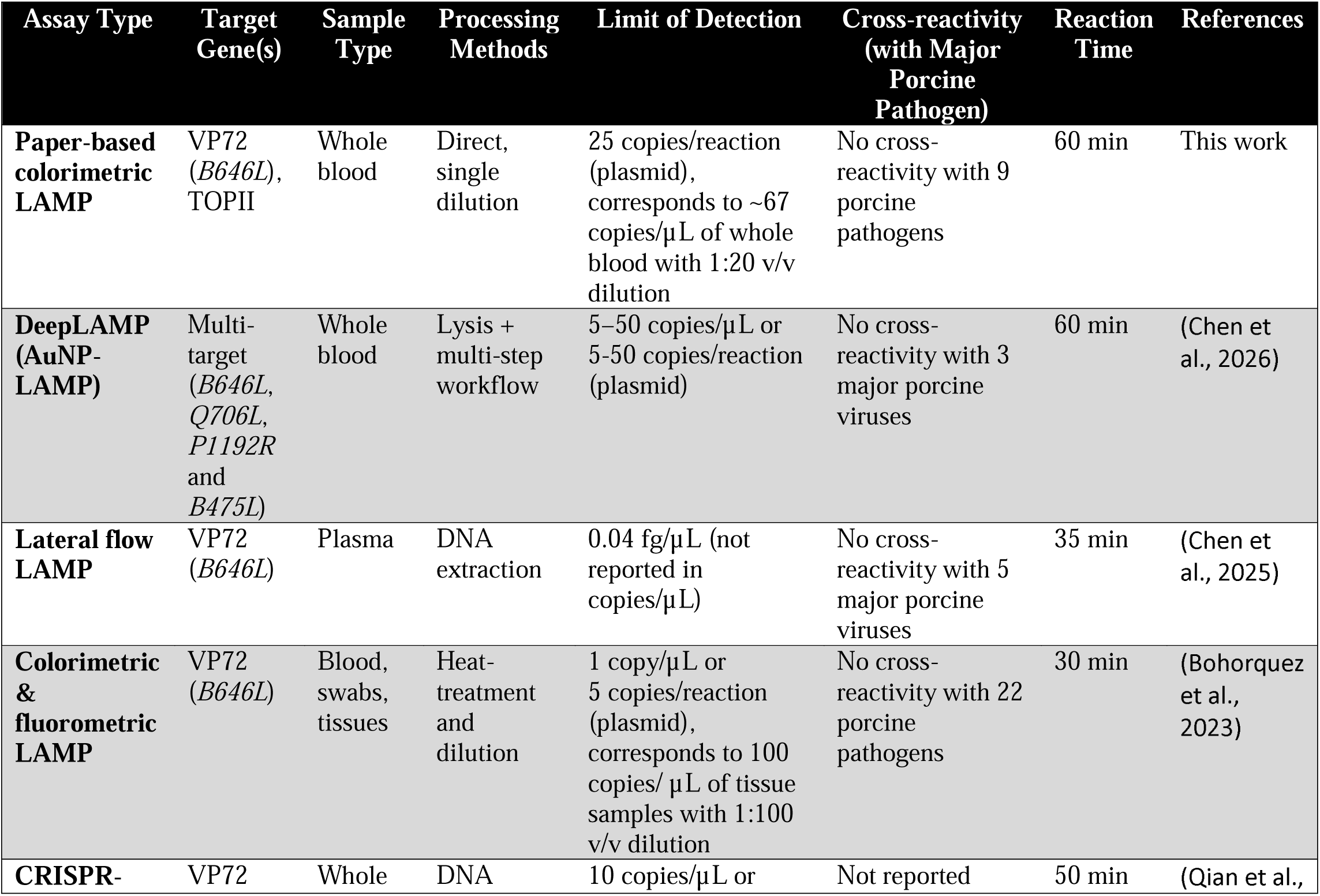
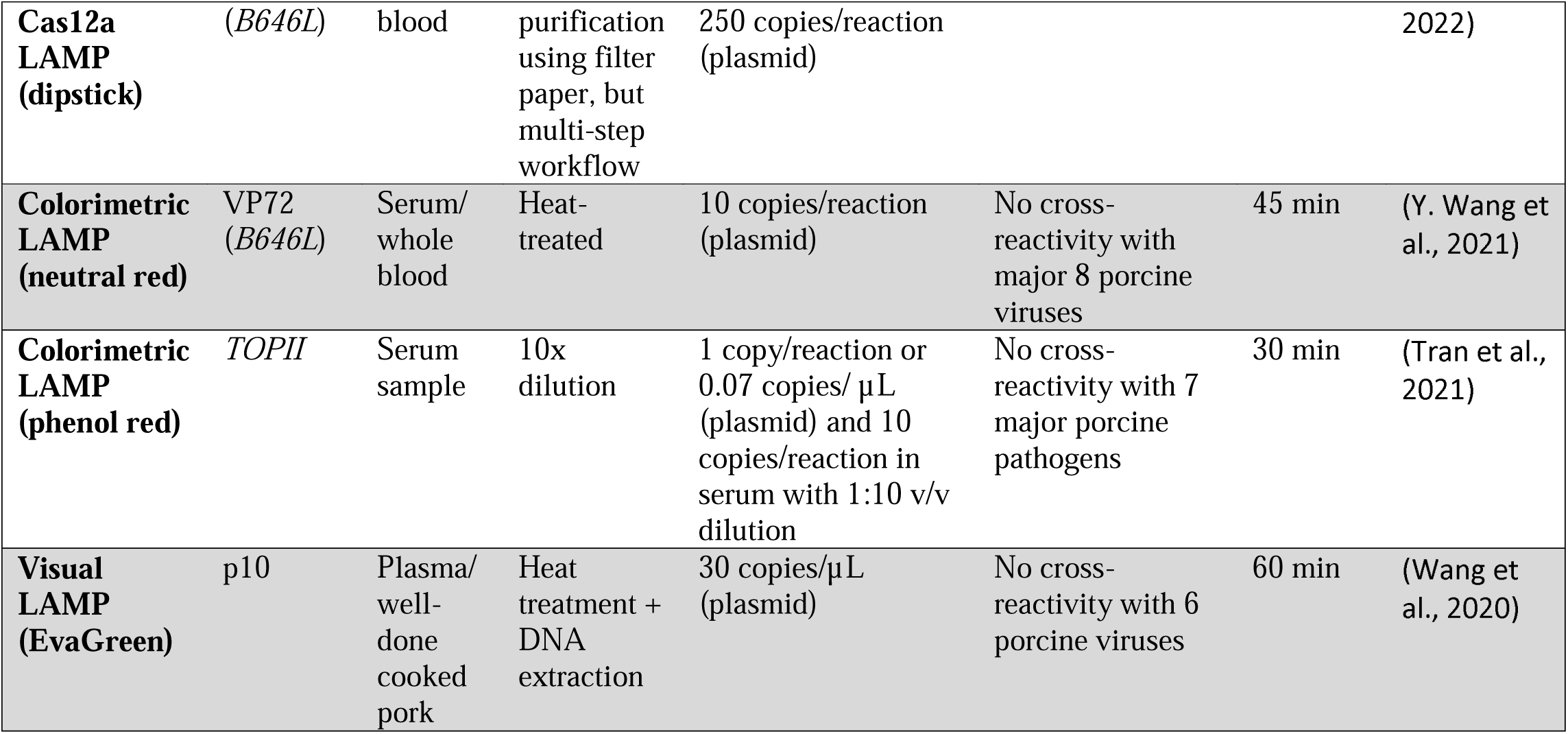
Point of need (PON) molecular DNA tests for African swine fever virus (ASFV) detection based on loop-mediated isothermal amplification (LAMP) assays reported between 2020 and 2026.

Despite these advances in PON molecular platforms, whole blood remains a challenging matrix for one-pot nucleic acid amplification because blood contains amplification inhibitors and exhibits strong buffering capacity (Hu et al., 2023; Nwe et al., 2024). This issue is particularly problematic for colorimetric LAMP assays that rely on pH-sensitive dyes such as phenol red, as the high buffering capacity of blood can suppress amplification-driven pH changes or mask out colorimetric signals (Gu et al., 2025; Raut et al., 2026a; Sritong et al., 2023; Zhang et al., 2025). Consequently, many reported molecular workflows continue to rely on nucleic acid extraction, preheating, sample dilution, or other preprocessing steps, effectively converting field testing into a multi-step (typically 3–6 step) sample preparation process that often requires additional reagents and customized instrumentation with off-board analysis for reliable data acquisition and interpretation.

Here, we report a portable, PON molecular diagnostic platform for ASFV detection that enables colorimetric qLAMP directly from diluted whole blood on microfluidic paper-based analytical devices (µPADs) (Fig. 1). We targeted the conserved VP72 capsid protein (*B646L* gene) and *TOPII* genes and designed the platform to eliminate reliance on nucleic acid extraction and purification while reducing subjective visual interpretation through on-board objective colorimetric signal analysis. Using plasmid DNA spiked into whole blood diluted to 5% (v/v) in 5% D-mannitol, the assay achieved a LOD of 25 copies per reaction (67 copies/µL of blood sample) for the VP72 target with no cross-reactivity against 9 major porcine viral and bacterial pathogens, demonstrating 100% analytical sensitivity and specificity during in-house testing (Fig. 1B-C) and 90% and 92% analytical sensitivity and specificity respectively in an external laboratory evaluation (Fig. 1D-E). The complete assay, including isothermal incubation and automated result reporting, was completed within 60 min using a custom-built portable heating and imaging system (benchtop ThermiQuant™ MegaScan (Raut et al., 2026a) and portable ThermiQuant™ AquaStream (Raut et al., 2026c) with onboard software). Together, these results demonstrate a simplified, extraction-free molecular diagnostic approach that integrates paper-based assay chemistry with objective signal interpretation to support rapid and reliable ASFV detection at the PON.

**Fig. 1.**
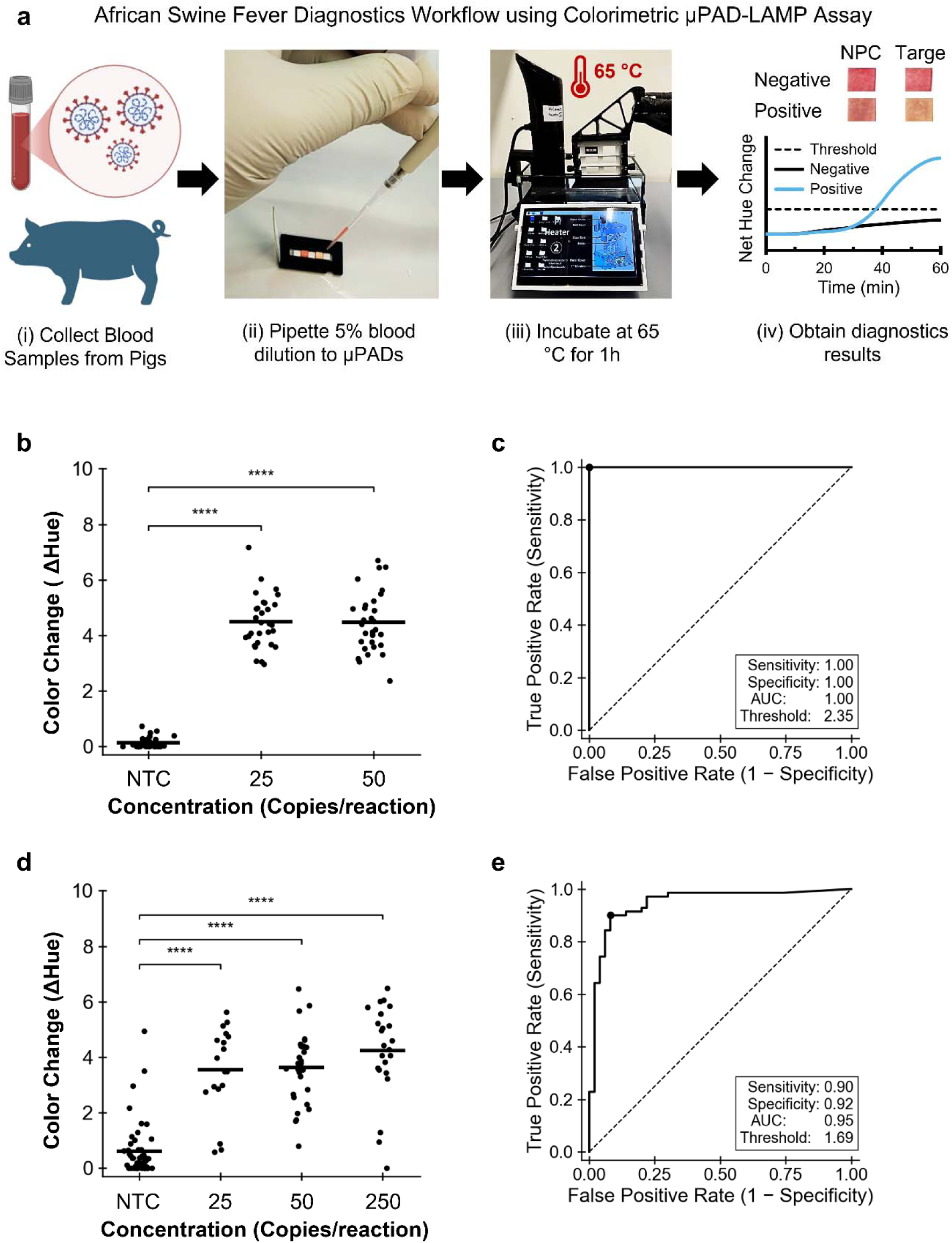
Point of need (PON) workflow and inter-laboratory performance of the µPAD-LAMP assay. (a) Field-ready µPAD-LAMP workflow showing (i) blood collection in EDTA vacutainers followed by dilution to 5% (v/v) in 5% D-mannitol, (ii) positive and negative samples pipetting onto no primer control (NPC) and target µPADs followed by cartridge sealing using an integrated sealing tape, (iii) incubation in the ThermiQuant™ AquaStream portable water-bath heater at 65 °C, and (iv) result output from the portable instrument as both a visible color change and real-time hue–time plots with automatic thresholding to classify positive and negative reactions. (b) Net hue changes distributions (swarm plot) obtained from in-laboratory testing using pooled pig blood diluted to 5% in D-mannitol and spiked with VP72 plasmid DNA at 0 copies per reaction (no-template control, NTC), 1× the limit of detection (LOD; 25 copies per reaction), and 2× LOD (50 copies per reaction). (c) Receiver operating characteristic (ROC) curve corresponding to in-laboratory testing shown in (b), demonstrating 100% analytical sensitivity and 100% analytical specificity. (d) Net hue changes distributions obtained from an external collaborative laboratory using the same workflow with expanded concentration levels (0, 1×, 2×, and 10× LOD). (e) ROC curve corresponding to external laboratory testing shown in (d), yielding 90% analytical sensitivity and 92% analytical specificity.

## Methods

### ASFV synthetic plasmid targets synthesis and quantification

Synthetic DNA plasmids containing TOPII (NCBI: Z14245.1) and VP72 (NCBI: MG209614.1) were obtained from GenScript Biotech Corporation, USA and quantified by digital PCR (dPCR). dPCR primer and probe sequences are listed in Table S1, and template sequences are provided in the Supporting Information (SI; VP72_TOPII_Template_Sequence.xlsx”). Additional methodological details are provided in SI Note 1.1.

### ASFV LAMP primer design, synthesis, and screening

Six new LAMP primer sets were designed for each target gene (VP72 and TOPII) using PrimerExplorer V5 (Eiken Chemical Co., Ltd.). One previously published primer set each for *TOPII* (James et al., 2010) and *B646L* (VP72) (Y. Wang et al., 2021) was included as a reference and designated as TOPII.1 and VP72.1, respectively. All primer sequences are provided in Table S2, and the final primer set (VP72.4) used for paper LAMP validation is listed in Table 2. For primer screening, fluorescent qLAMP reactions were performed using WarmStart® 2× LAMP Master Mix (New England Biolabs, USA) in 5-µL reaction volumes and monitored in real time at 65 °C for 60 min using a qTower 384 system (Analytic Jena, Germany). dPCR-quantified templates were serially diluted, and no-template controls (NTCs) were included as negative controls, with fluorescence signals collected in the FAM channel. Primer sets were ranked using five performance metrics implemented in the Primer Scoring software (Kamel et al., 2025b; Pascual-Garrigos et al., 2021). Additional experimental details are provided in SI Note 1.2.

**Table 2.**
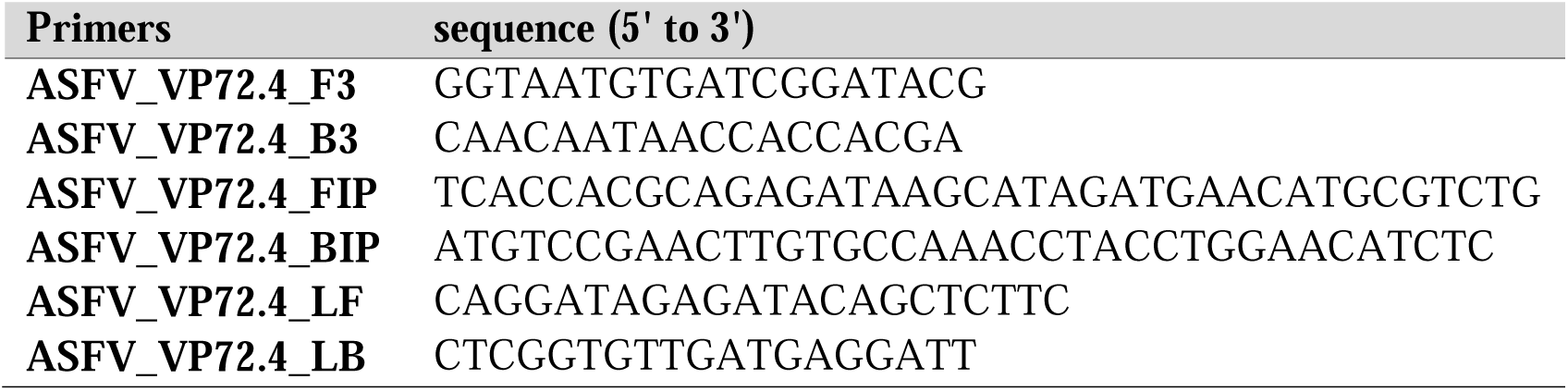
Nucleotide sequences of final LAMP primer set used for paper LAMP.

### µPAD and acrylic cartridge fabrication

µPADs (3 mm× 3 mm) were fabricated as previously described (Ahmed et al., 2025; Raut et al., 2026a). Briefly, grade 222 chromatography paper (Ahlstrom-Munksjö, Finland) and polystyrene spacer sheets (HIPS Litho Grade, Tekra, USA) were cut into 3-mm strips, assembled onto adhesive-backed polyester film (MELINEX® 454, Tekra, USA), and sectioned into two-, three-, or four-pad µPAD strips. Acrylic cartridges for housing the µPAD strips were fabricated following previously reported designs (Raut et al., 2026a, 2026c), designed in SolidWorks (Dassault Systèmes, France), and laser-cut from 1.5-mm acrylic sheets. Laser fabricated acrylic cartridges were sequentially wiped and then dried, with reverse osmosis (RO) water, 70% ethanol, and RNase AWAY™ prior to use. The cartridge design was adaptable to different µPAD configurations and slot geometries. Additional details are available in SI Note 1.3.

### Preparation of colorimetric paper LAMP and assembly of µPADs into cartridge

The colorimetric LAMP formulation for the paper-based assay was adapted from previously reported SARS-CoV-2 detection protocols (Davidson et al., 2021; J. Wang et al., 2021b). Briefly, a homemade 2× LAMP master mix containing buffer components, deoxynucleoside triphosphate (dNTPs)/ deoxyuridine triphosphate (dUTP), Antarctic Thermolabile Uracil-DNA Glycosylase (UDG), Bacillus stearothermophilus (Bst) 2.0 DNA polymerase, phenol red, and stabilizing additives were prepared (full composition provided in Table S3). The final reaction mixture included target-specific primers, additional polymerase, betaine, BSA, and trehalose (final concentrations summarized in Table S4). Aliquots (7.5 µL) were dispensed onto µPADs, air-dried (∼2 h), assembled into acrylic cartridges, sealed with PCR film, and used immediately or stored at −20 °C for up to one week. Additional details are in SI Note 1.4.

### Cross-reactivity against other major swine pathogens

Nine major swine pathogens were used for cross-reactivity testing using ASFV VP72.4 primer set. These included five bacteria (*Actinobacillus pleuropneumoniae*, *Actinobacillus suis*, *Bordetella bronchiseptica*, *Pasteurella multocida*, and *Streptococcus suis*) and four viruses: Influenza A (H1N1, swine-origin strain VR-1682), Porcine Respiratory Coronavirus (PRCV, strain VR-2384), Porcine circovirus type 2d (PCV2d), and Porcine Reproductive and Respiratory Syndrome Virus 2 (PRRSV2, strain: VR-2332). All bacterial pathogens were cultured, and genomic DNA was extracted using the Invitrogen PureLink™ gDNA Kit. Synthesized genomic DNA for PCV2d and PRRSV2 was acquired from GenScript Biotech Corporation in the pUC57 plasmid, while PRCV and H1N1 viral strains were procured from ATCC (USA) and cultured according to ATCC protocols, after which viral RNA was extracted using the Invitrogen PureLink™ Viral RNA/DNA Kit. All pathogen targets were quantified using dPCR, and the corresponding primers and probes were obtained from published sources and are listed in Table S1.

Whole blood was collected from adult pigs into EDTA-coated vacutainer tubes. All animal procedures were approved by the Purdue University Institutional Animal Care and Use Committee (IACUC number: IPROTO2205002265). For cross-reactivity test, plasma was used and was separated by centrifugation at 1,500 × g for 10 min at room temperature and subsequently diluted to 5% (v/v) in reverse osmosis (RO) water. The diluted plasma samples were spiked with the ASFV VP72 target and the nine swine pathogens described above at a concentration of 10^3^ copies per reaction. Reactions included no-primer controls (NPCs) and NTCs. In addition, a VP72-spiked positive control prepared in nuclease-free water without plasma was included for comparison.

### Optimization of isotonic diluents for whole blood–based colorimetric paper LAMP assay

To identify an isotonic blood diluent that minimizes the inhibitory effects of heme and other blood components on colorimetric LAMP amplification, whole blood was diluted to a final concentration of 5% (v/v) using a panel of candidate diluents.

Seven diluents were prepared in nuclease-free water: 0.9% (w/v) NaCl, 5% (w/v) D-glucose, 5% (w/v) D-fructose, 5% (w/v) glucose, 5% (w/v) D-myo-inositol, 5% (w/v) D-mannitol, and 10% (w/v) sucrose. All chemicals used for diluent preparation were of analytical grade. NaCl was purchased from Fisher Chemical (Thermo Fisher Scientific, USA). D-glucose and sucrose were obtained from Alfa Chemistry (USA), while D-fructose, D-myo-inositol, and D-mannitol were purchased from Sigma-Aldrich (USA). Stock solutions of each diluent were prepared in advance and stored at room temperature, and whole blood dilutions were freshly prepared immediately before each experiment. For assay evaluation, VP72 plasmid DNA was spiked into diluted whole blood and colorimetric paper LAMP reactions were performed using the VP72.4 primer set. To assess the effect of blood concentration on assay performance, whole blood was diluted to final concentrations of 5%, 10%, 20%, 40%, 60%, 80%, and 100% (v/v) using 5% (w/v) D-mannitol as the diluent. Each dilution was spiked with VP72 plasmid DNA, and colorimetric paper LAMP reactions were conducted using the VP72.4 primer set.

### Limit of detection and limit of quantification of the paper-based colorimetric ASFV LAMP assay

To determine the LOD of the paper-based colorimetric LAMP assay, whole blood was diluted to 5% (v/v) in 5% (w/v) D-mannitol and spiked with serially diluted concentrations of the VP72 plasmid target, with NTCs included as negative controls. µPAD-LAMP reactions were performed with 3–6 technical replicates per concentration. The initial LOD was determined using probit analysis (Klymus et al., 2020; Stokdyk et al., 2016a) and the LOD95 was defined as the lowest analytical concentration predicted to yield a positive detection in 95% of reactions in a serially diluted series with three technical replicates per concentration. The final LOD was then confirmed using the standard concentration closest to the predicted LOD95 (25 copies per reaction in this study). Twenty technical replicates at LOD95 were tested at this concentration, along with twenty negative controls. The final LOD was considered when at least 19 of 20 of spiked replicates showed positive amplification, while all negative controls remained non-detected.

The limit of quantification (LOQ) was defined as the lowest DNA concentration for which the coefficient of variation (CV) of the quantification time (Tq) was ≤10% (CV = 100 × (SD/Mean) and the linear regression on the standard calibration curve exhibited strong linearity (R^2^ > 0.9).

To further evaluate the analytical sensitivity and specificity of the assay (in house testing), three target concentrations (1×, 2×, and 0× (NTC) of LOD) were tested with 30 technical replicates each. A similar experiment was conducted at an external collaborating laboratory using an expanded concentration range (1×, 2×, 10×, and 0× (NTC) of LOD). For the external evaluation, a written user manual (SI, “ASF_Field_Manual.docx”) was provided along with whole blood samples, prepared µPADs preloaded with dried LAMP reagents, and target templates. In addition, authors GP and NR conducted a single on-site visit to the external laboratories to provide hands-on demonstration and training.

## Results

### Field-ready ASFV workflow enables robust inter-laboratory performance of the µPAD-LAMP assay

We developed a field-deployable colorimetric µPAD-LAMP diagnostic kit that integrates sample handling, amplification, and automated result interpretation into a single PON workflow for the detection of ASFV (Fig. 1A). In this workflow, pig blood collected in EDTA vacutainer was diluted to 5% (v/v) in 5% D-mannitol and directly pipetted to µPADs containing pre-dried colorimetric LAMP reagents housed within an acrylic cartridge. After sample addition (VP72 target, 7.5 µL per reaction zone), the cartridge was sealed with foldable PCR tape that created a watertight, evaporation-resistant environment for 65°C incubation. Up to six sealed cartridges were assembled simultaneously in a custom portable water-bath incubator (ThermiQuant™ AquaStream (Raut et al., 2026c)) maintained at 65 °C and equipped with an onboard camera and Raspberry Pi–based software for automated time-lapse imaging and real-time colorimetric analysis. During incubation, the system continuously extracted hue information (every 30 seconds) and displayed LAMP amplification trajectories, enabling both visual endpoint interpretation and objective, software-based result determination. At the conclusion of the 60-min reaction time, AquaStream provided automated result output along with access to raw and processed data, enabling standardized and objective PON ASFV detection following manual sample loading.

Using this field-ready workflow, we evaluated the performance of the µPAD-LAMP assay under conditions designed to approximate PON deployment. Because ASFV is not present in the United States, we assessed diagnostic performance using pooled pig blood samples spiked with defined concentrations of VP72 plasmid DNA rather than conducting field trials with clinical specimens. We performed this evaluation both in our laboratory, with operators familiar with the µPAD-LAMP workflow, and in an external collaborative laboratory with operators experienced in molecular assay handling but unfamiliar with the paper-based format and given minimal training. Fig. 1B-C show results obtained in our laboratory. Fig. 1B presents net hue change distributions for pooled pig blood samples diluted to 5% in D-mannitol and spiked with VP72 plasmid DNA at 0 copies per reaction (NTC), 1×LOD (25 copies per reaction), and 2× LOD (50 copies per reaction). Positive samples exhibited a clear and statistically significant increase in ne hue change relative to negative controls, and receiver operating characteristic (ROC) analysis Fig. 1C) demonstrated 100% analytical sensitivity (60/60) and 100% analytical specificity (30/30).

Fig. 1D-E show results obtained in the external laboratory using the same workflow but expanded concentration levels (0, 1×, 2×, and 10× the LOD). Fig. 1D shows the corresponding net hue change distributions. Although positive samples remained distinguishable from negative controls, increased overlap between groups was observed relative to in-laboratory testing (Fig. 1B). ROC analysis of this data (Fig. 1E) yielded an analytical sensitivity of 90% (63/70) and a specificity of 92% (46/50), reflecting reduced but still high analytical performance. This decrease in performance likely resulted from a combination of differences in handling of the paper-based format and extended storage and transport times of the paper biosensors prior to testing, including longer test durations (one week versus two days for external lab and in-lab testing respectively) and repeated freeze–thaw cycles of the synthetic templates. Despite this reduction, the assay maintained greater than 90% analytical sensitivity and specificity in an external laboratory setting, supporting the robustness of the µPAD-LAMP assay for PON deployment. Further validation using clinical samples in field settings will be required to fully establish clinically relevant diagnostic performance.

Moreover, the cost of goods (COG) of the µPAD-assay including the cost of air tight cartridge was USD 1.69 per single test that included 1 control and 1 test µPADs (Table S7).

### Primer screening results

The ASFV *B646L* gene, encoding the VP72 (p72) major capsid protein, and the viral *TOPII* gene, which is involved in ASFV DNA replication, were selected as diagnostic targets for ASFV detection based on their high sequence conservation across ASFV isolates (Hu et al., 2023; Jia et al., 2017; Lim et al., 2023; Zhang et al., 2025). The *B646L* gene is the primary molecular target recommended by the WOAH for ASFV diagnostics (WOAH, 2024), while the *TOPII* gene was selected based on its previously demonstrated utility in ASFV detection assays (Tran et al., 2021).

We screened 14 LAMP primer sets (seven targeting TOPII and seven targeting VP72) using fluorescence-based qLAMP with serially diluted synthetic plasmid templates ranging from 10^4^ to 0 (NTC) copy per reaction. Fig. S1 (TOPII) and Fig. S2 (VP72) show the fluorescence vs time curves for all primer sets and Fig. 2 show representative set at 10^3^ copies/reaction. We evaluated primer performance using the previously published Primer Scoring tool (Kamel et al., 2025b; Pascual-Garrigos et al., 2021), and Tables S5 and Table S6 summarize the ranking results for TOPII and VP72 primer sets respectively. In both target regions, our newly designed primer sets outperformed the published reference primers (TOPII.1 and VP72.1). Among them, VP72.4 showed the best overall performance (rank1, overall score: 100/100), and was therefore selected for subsequent optimization in µPAD-LAMP assay.

**Fig. 2.**
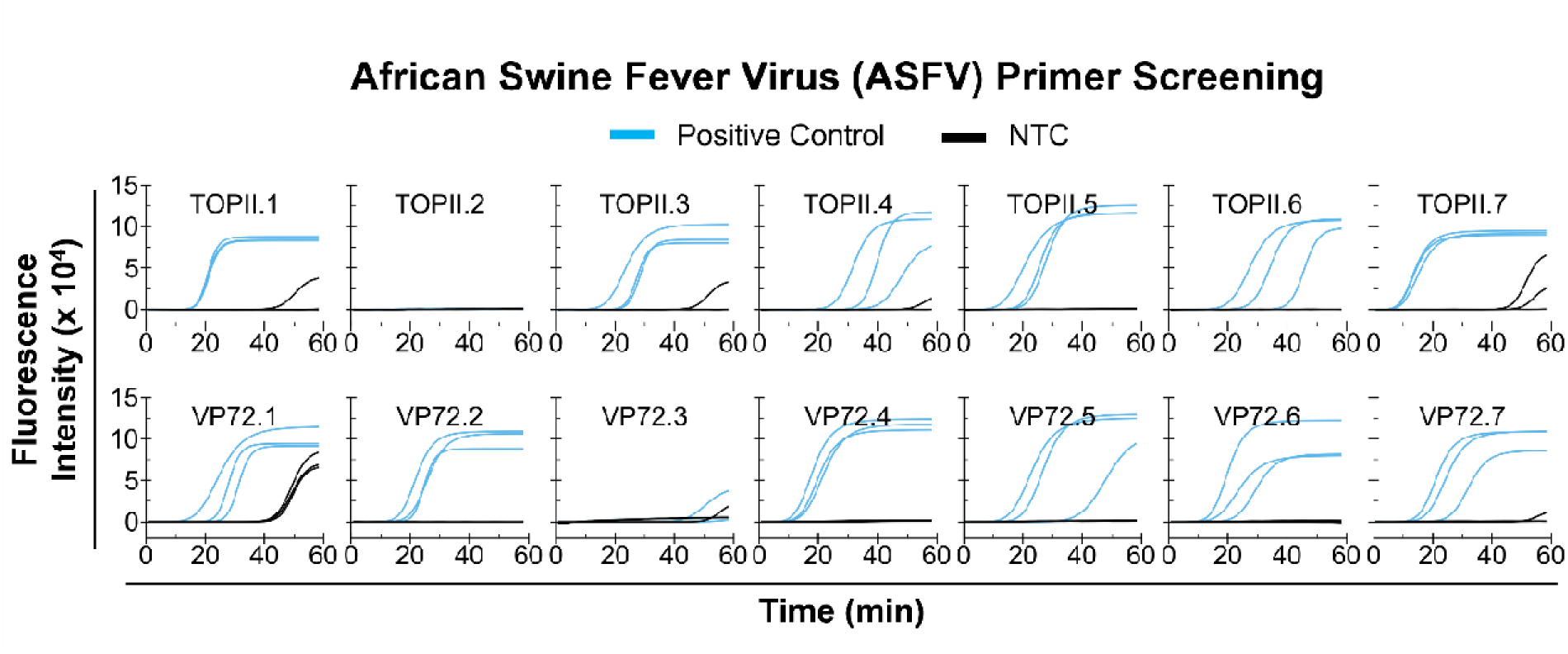
Screening of ASFV LAMP primer sets. Fluorescence vs. time curves of primers targeting VP72 and TOPII of African swine fever virus (ASFV) obtained using a fluorescent LAMP assay with the NEB WarmStart® DNA/RNA LAMP Kit and synthetic DNA templates.

### µPAD-LAMP biosensor achieves a 25-copies/reaction limit of detection (LOD) in diluted whole blood

We repurposed our previously reported µPAD strip design with alternating µPAD and spacer to enable parallel reactions while preventing fluidic cross-talk between adjacent reaction zones (Fig. 3A) (Ahmed et al., 2025; Davidson et al., 2021; Kamel et al., 2025b; Raut et al., 2026a). Each µPAD contained preloaded, dried colorimetric LAMP reagents that rapidly (within seconds) rehydrated upon sample addition. We housed the assembled µPAD strips within an airtight acrylic cartridge (1.5 mm thick), which prevented reagent and sample vapor leakage during the 1h incubation at 65 °C. Upon amplification, positive reactions produced a red-to-yellow color transition, whereas negative reactions remained red (Fig. 3A, iv), providing both a visual readout and a signal that could be captured using a scanner or camera and quantified by digital hue extraction for objective interpretation (Ahmed et al., 2025; Raut et al., 2026a, 2026c; Wang et al., 2024).

**Fig. 3.**
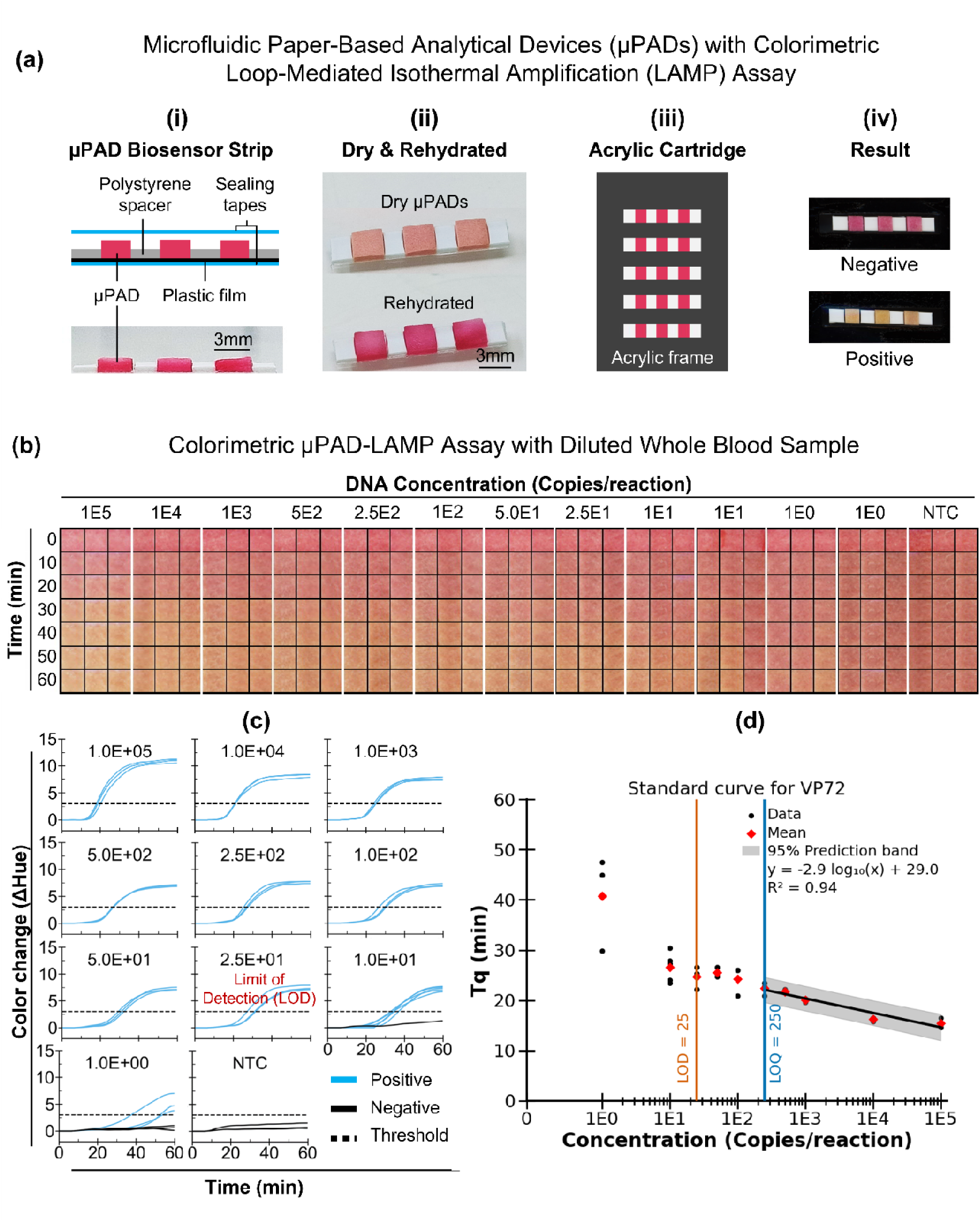
African swine fever virus (VP72 target) detection in diluted whole blood using a paper-based colorimetric LAMP assay. (a) Schematic of the µPAD-LAMP biosensor showing (i) exploded view of µPAD–spacer architecture used to prevent crosstalk between adjacent µPADs, (ii) µPADs after reagent drying and rehydration, (iii) housing of µPAD strips within an airtight acrylic cartridge to prevent evaporation, and (iv) representative positive (yellow) and negative (red) colorimetric results. (b) Time-stamped cropped images (0–60 min, 10-min intervals) of colorimetric µPAD-LAMP reactions containing 5% adult swine blood diluted in 5% D-mannitol and spiked with VP72 plasmid DNA (10^5^ to 1 copy per reaction; n = 3, except 10 and 1 copies/reaction where n = 6), along with no-template controls (NTCs; n = 3). (c) Net-hue–versus–time trajectories extracted from timelapse images shown in (b). (d) Standard calibration curve showing quantification time (Tq) plotted against the log_10_ (VP72 template concentration).

Using the µPAD strip design we evaluated the analytical sensitivity of the colorimetric µPAD-LAMP sensor using EDTA whole blood diluted to 5% (v/v) in 5% D-mannitol and spiked with VP72 plasmid targets ranging from 10^5^ to 0 (NTC) copies per reaction. Fig. 3B shows cropped reaction-zone images captured using our custom-built ThermiQuant™ MegaScan heater-imager instrument (Raut et al., 2026a), with representative images from 0 to 60 min at 10-min intervals (images displayed with +20% contrast and +40% brightness). Positive reactions exhibited a color change from red to yellow, whereas negative reactions remained red. Fig. 3C shows the corresponding net-hue-versus-time profiles extracted using our custom image analysis software (Amplimetrics™ (Raut et al., 2026a)). We applied a positivity threshold of 3 hue units to distinguish sigmoidal amplification trajectories from linear negative responses. Given the limited replicate number (n = 3 or 6), we defined preliminary 95% limit of detection (LOD95) as the lowest target concentration that crossed the 95% detection frequencies in a probit analysis (Stokdyk et al., 2016b) plot and obtained LOD95 as 26 copies/reaction (Fig. S3A). For final LOD, we used 20 technical replicates near the LOD95 (at 25 copies/reaction) and 20 NTCs and confirmed the final LOD as the lowest concentration that yielded successful amplification in at least 19/20 positive replicates and none in NTCs. This final LOD of 25 copies/reaction corresponds to 3.3 copies/µL in the 5% diluted blood sample used in the assay, which is equivalent to 67 copies/µL in the original whole blood sample after accounting for the 20-fold dilution.

We defined the LOQ as the lowest standard concentration with a CV ≤10%. LAMP assays generally exhibit low variability, with reported CVs typically ∼0–10% and occasionally up to ∼15% (Hayes et al., 2025; Lalonde et al., 2021; Lou et al., 2024; Wu et al., 2022). Although ASFV LAMP studies do not consistently report CV, available data from liquid assays show low variability (∼1–6%) under controlled conditions (Ji et al., 2023; Mee et al., 2020; Wang et al., 2022). Given that our assay uses a paper-based format with dried reagents, we adopted a slightly relaxed threshold and selected a 10% CV as a practical criterion for quantification. This threshold differs from qPCR, where substantially higher CV cutoffs (up to ∼35%) have been reported (Klymus et al., 2020). Using the 10% CV threshold, we determined the LOQ to be 250 copies per reaction (660 copies/µL of whole blood) (Fig. S3B). Fig. 3C shows a standard calibration curve describing the relationship between the quantification time (Tq), defined as the time corresponding to the peak of the second derivative of the hue–time curve (Fig. 3B), and the log_10_-transformed VP72 template concentration (copies per reaction). The Tq values decreased with increasing template concentration and exhibited a strong linear relationship over the tested range of 10^5^ to LOQ (250 copies/reaction), with an R^2^ of 0.94, demonstrating the quantitative capability of the µPAD-LAMP biosensor.

We further evaluated the analytical sensitivity of the WOAH-recommended King’s qPCR assay (King et al., 2003) and the NAHLN qPCR assay (with minor sequence modifications; see Table S1, target: ASFV_VP72 (Set 2) (Zsak et al., 2005)) using serially diluted VP72 targets spiked either into 5% D-mannitol or into whole blood diluted to 5% in D-mannitol, corresponding to a final blood concentration of 1% (v/v) in the qPCR reaction. Fig. S4A–B show the corresponding normalized fluorescence–time amplification curves for the King’s and NAHLN assays, and Fig. S4C summarizes the LOD and LOQ for each assay. We determined the LOD using probit analysis and the LOQ using a 10% CV threshold. Both qPCR assays achieved an LOD of 4 copies/reaction and an LOQ of 10 copies/reaction in non-blood samples. In blood samples, performance decreased slightly, with LODs of 11 and 13 copies/reaction for the King’s and NAHLN assays, respectively, while the LOQ was 25 copies. The qPCR assays showed approximately two-fold better LOD than the paper LAMP assay, indicating comparable sensitivity. However, the paper LAMP assay exhibited higher variability, resulting in an approximately ten-fold worse LOQ than the qPCR assays.

### µPAD-LAMP biosensor shows high specificity against common swine pathogens

We evaluated specificity of the final selected primer set (VP72.4) against nine common swine pathogens using the colorimetric µPAD-LAMP biosensor with EDTA plasma samples diluted to 5% in water. Plasma was used in this study because the experiments were conducted prior to optimization of the dilution protocol for whole blood samples. Fig. 4A shows the colorimetric responses of the µPAD-LAMP biosensor at 60 min, while Fig. 4B presents the corresponding hue–time profiles for each target. The VP72.4 primer set produced a positive signal only in the presence of African swine fever virus VP72 targets, either spiked into 5% plasma or tested in water, and showed no detectable amplification with other swine pathogens. In addition, neither NPC nor NTC produced false amplification, demonstrating the high specificity of the assay toward clinically relevant swine pathogens.

**Fig. 4.**
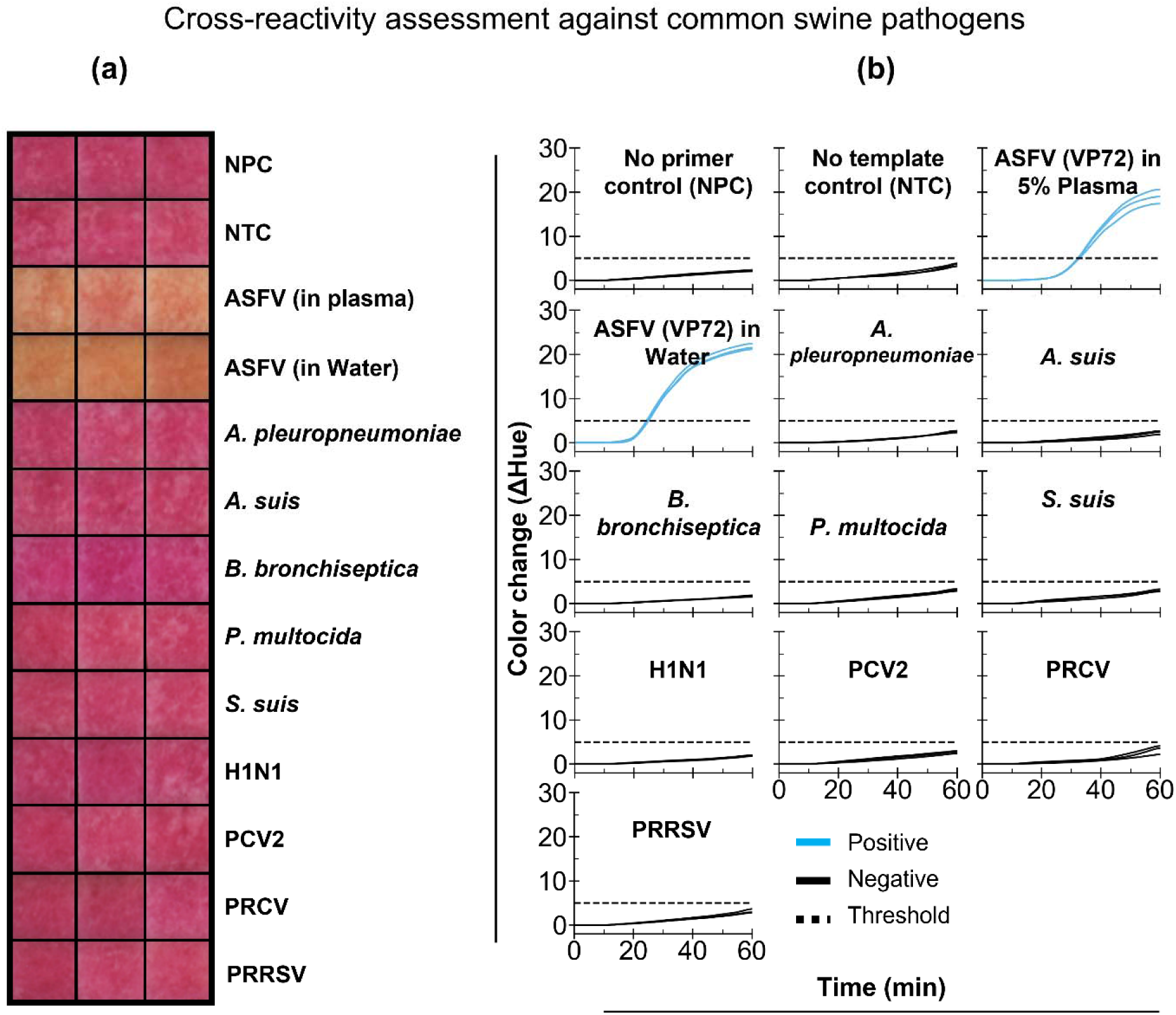
Cross-reactivity assessment of the paper-based loop-mediated isothermal amplification (LAMP) biosensor for African swine fever virus (ASFV) detection targeting the VP72 gene. (a) Colorimetric µPAD-LAMP results at 60 min using samples spiked with the ASFV VP72 target in water, the ASFV VP72 target in 5% diluted plasma, and other major swine pathogen targets spiked in 5% diluted plasma, including *Actinobacillus pleuropneumoniae*, *Actinobacillus suis*, *Bordetella bronchiseptica*, *Pasteurella multocida*, *Streptococcus suis*, swine influenza A virus (H1N1), porcine circovirus type 2d (PCV2d), porcine respiratory coronavirus (PRCV), and porcine reproductive and respiratory syndrome virus 2 (PRRSV2, strain: VR-2332). A no-primer control (NPC; VP72 target without primers) and a no-template control (NTC) were used as negative controls. Images were adjusted (−20% brightness, +40% contrast) in Microsoft PowerPoint for visualization. (b) Corresponding hue–time plots generated from the time-lapse images shown in (a), illustrating positive and negative amplification responses.

### µPAD-LAMP biosensors are robust to blood sample age and inter-animal variability but require dilution

To evaluate the impact of whole-blood matrix effects on paper-based colorimetric LAMP performance, we first examined the effect of blood concentration on assay readout. We tested whole blood (3 weeks old, stored at 4 °C) diluted from 5% to 100% in 5% D-mannitol and spiked with 10^5^ copies/reaction of VP72 plasmid DNA. Fig. 5A shows cropped reaction-zone images acquired at 0, 10, and 60 min across the tested blood dilutions. Increasing blood concentration caused the paper substrate to appear progressively darker red at 0 min. After incubation at 65 °C, samples containing >5% blood turned brown and failed to develop the characteristic yellow color indicative of successful amplification, whereas only the 5% blood dilution produced a clear red-to-yellow transition. Based on these results, we selected 5% whole blood as the working dilution for subsequent experiments.

**Fig. 5.**
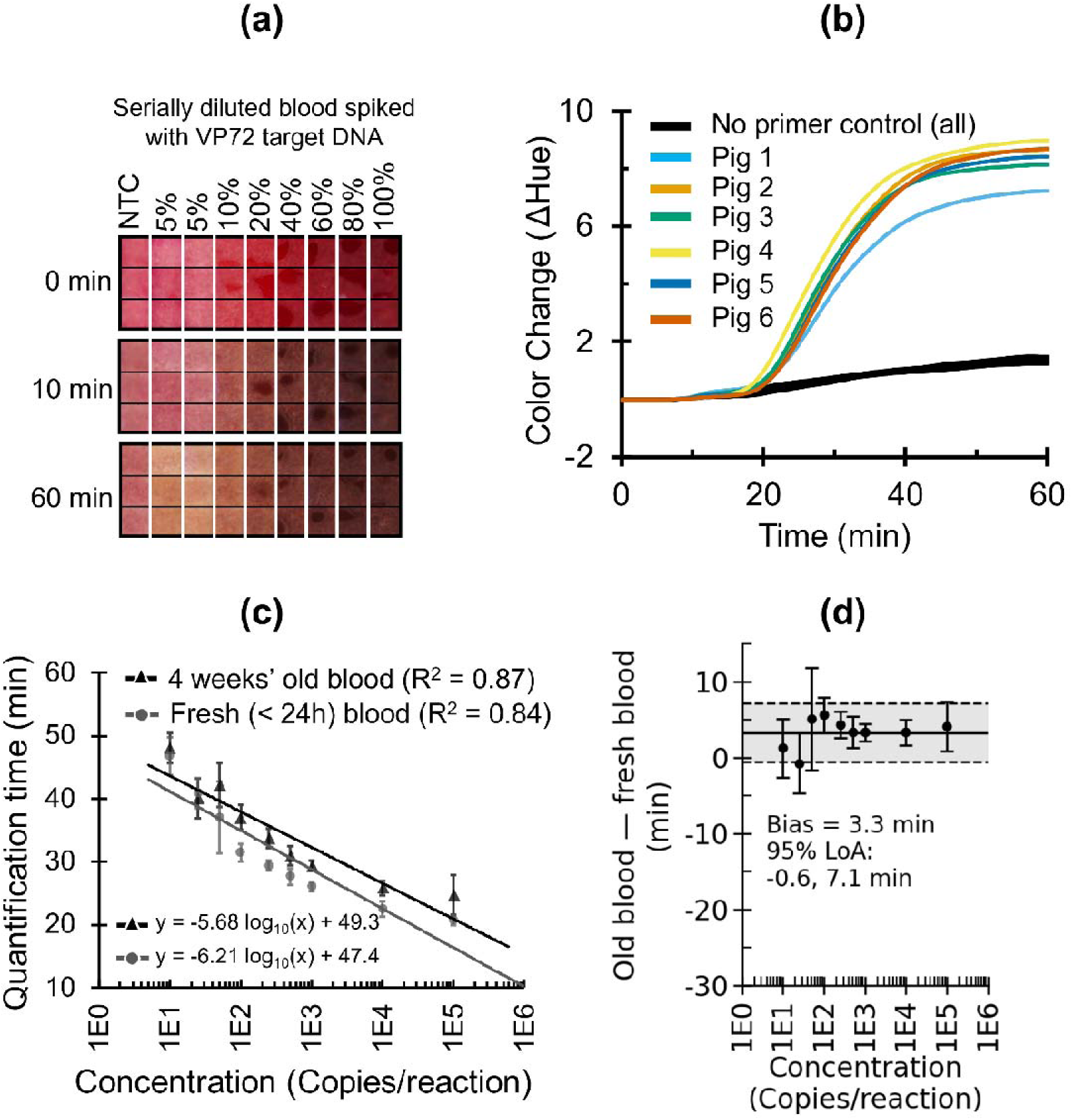
Effect of blood dilution, inter-animal variability, and blood age on paper-based colorimetric LAMP performance. (a) Cropped reaction-zone images acquired at 0, 10, and 60 min from µPAD-LAMP assays containing serial whole-blood dilutions (5–100% v/v) prepared in 5% D-mannitol and spiked with VP72 synthetic target or without (no template control). (b) Net hue–versus–time trajectories showing colorimetric responses for blood samples obtained from six different animals spiked with 10^5^ copies/reaction of VP72 and no template control. (c) Standard calibration curves showing quantification time (Tq) plotted against the log - transformed VP72 template concentrations (25 to 10^5^ copies/reaction) for fresh (<24 h) and aged (4 weeks old) whole blood diluted to 5% (v/v) in 5% D-mannitol. (d) Bland–Altman plot comparing Tq values from fresh and aged blood in (c).

We evaluated whether inter-animal variability affected the performance of the µPAD-LAMP assay by testing fresh whole blood samples collected from six different adult pigs. We diluted each blood sample to 5% (in 5% D-mannitol) and spiked it with 10^5^ copies/reaction of VP72 plasmid DNA, then monitored amplification using the colorimetric readout. Fig. 5B shows the corresponding hue–time profiles for all positive six pig blood samples and NPC for all. The assay produced successful amplification with comparable hue–time trajectories across all samples, indicating consistent µPAD-LAMP performance with blood obtained from different animals.

We next evaluated whether blood age influenced reaction kinetics by comparing fresh whole blood (<24 h old) and aged blood (4 weeks old, stored at 4 °C), each diluted to 5% in 5% D-mannitol. Fig. 5C shows standard calibration curves generated from concentrations at and above the LOD for both fresh and aged blood, which exhibit similar linear fits. Notably, we included all data points above the LOD, rather than restricting the analysis to concentrations above the LOQ, to enable comparison across a broader concentration range, including lower concentrations. Fig. 5D, a Bland–Altman analysis of the calibration data in Fig. 5C, shows that reactions prepared with fresh blood were, on average, 3.3 min faster than those prepared with aged blood; however, this difference represented less than 5% of the total Tq range, indicating that once diluted, blood age had a minimal impact on µPAD-LAMP assay performance under the tested conditions.

### µPAD-LAMP reactions containing 5% blood diluted in D-mannitol amplify faster than blood-free controls

Prior to selecting 5% D-mannitol as the optimal blood diluent, we screened isotonic solutions for compatibility with the colorimetric LAMP assay. We first evaluated representative ionic and non-ionic isotonic solutions by spiking each with 10^5^ copies/reaction of VP72 plasmid DNA. Fig. S5 shows endpoint (t = 60 min) cropped images of µPADs. The LAMP reaction failed to amplify in saline (0.9% NaCl), despite its widespread use as an isotonic solution. In contrast, reactions prepared in 5% D-mannitol and 5% D-glucose produced clear colorimetric amplification. Based on this initial screening, we focused subsequent experiments on non-ionic isotonic diluents, particularly carbohydrate-based formulations.

Next, we diluted pig whole blood stored at 4 °C for 27 days to 5% (v/v) in a panel of non-ionic isotonic solutions (5% D-fructose, 5% D-glucose, 5% D-myo-inositol, 5% D-mannitol, and 10% sucrose) or in water and compared these conditions with matching diluents without blood. We spiked all samples except NTCs with VP72 plasmid DNA at 10^5^ copies/reaction. Normalized hue–time curves (Fig. 6A) showed clear sigmoidal amplification trajectories for all spiked samples, whereas NTCs exhibited linear trajectories. However, two of three reactions containing blood diluted in water produced false-positive amplification, indicating that water may not be a suitable diluent for whole blood in colorimetric LAMP as water is hypotonic solution which causes hemolysis in red blood cells. Extracted Tq values for each condition are summarized in Fig. 6B. Among the isotonic conditions, 5% D-mannitol and 5% D-myo-inositol yielded the fastest amplification in blood-containing samples.

**Fig. 6.**
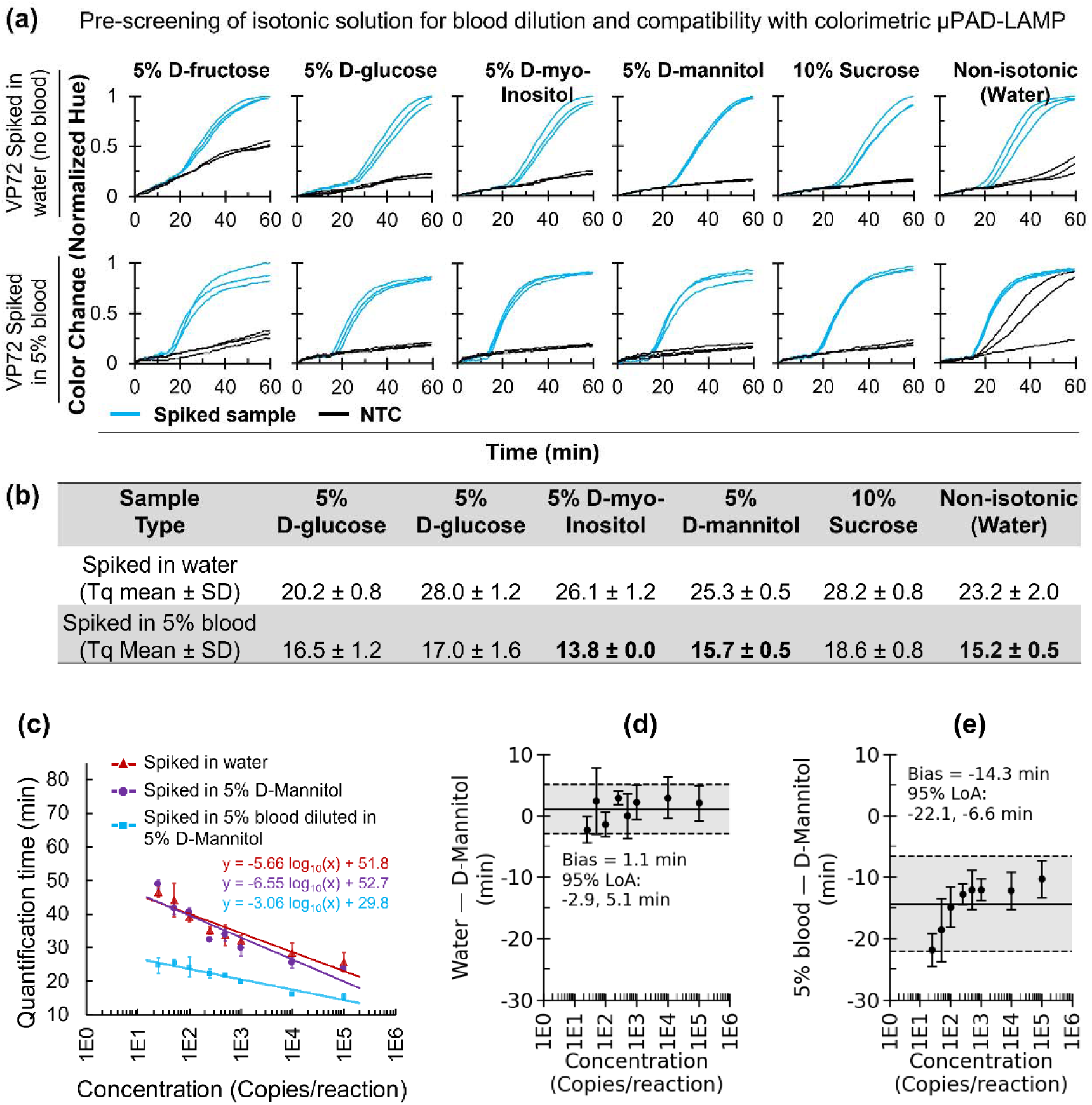
Effect of isotonic diluents and blood on colorimetric µPAD-LAMP amplification kinetics. (a) Normalized hue–versus–time trajectories showing pre-screening of isotonic solutions for blood dilution and compatibility with the colorimetric paper LAMP assay. Curves are shown for VP72 template spiked into water (baseline, top panel) and for 5% whole blood diluted with various isotonic solutions (bottom panel), along with the non-isotonic control (water alone). (b) Summary of quantification time (Tq) values extracted from (a), reported as mean ± SD. (c) Quantification time (Tq) plotted against VP72 template concentration for reactions prepared in water, 5% D-mannitol, or blood diluted to 5% in 5% D-mannitol. (d) Bland–Altman comparison of Tq values between LAMP reactions prepared in water and 5% D-mannitol. (e) Bland–Altman comparison of Tq values between LAMP reactions prepared with blood diluted to 5% in 5% D-mannitol and reactions prepared in 5% D-mannitol alone.

We therefore evaluated these two isotonic solutions further using serially diluted VP72 template (Fig. S6). Both diluents showed all technical replicates amplifying down to 10 copies per reaction (Fig. S6A, representing a 2.5-fold improvement over the previously observed LOD (Fig. 3B-C). Variability in LOD between runs on different day of experiments likely arose from differences in manual serial dilution and stochastic amplification within the paper matrix. Standard calibration curves, using all standards at or above LOD, showed strong log-linear relationships (R² = 0.84 for D-mannitol and 0.87 for D-myo-inositol; Fig. S6B), and Bland–Altman analysis (Fig. S6C) indicated close agreement between the two conditions (mean bias −1.0 min; 95% limits of agreement −5.4 to 3.4 min). Although the standard practice is to construct calibration curves using concentrations at and above the LOQ, we included all data above the LOD to enable comparison across a broader concentration range. Both diluents performed comparably and we selected 5% D-mannitol as the final diluent because of its wider use as a compatible osmolyte in diagnostic reagents.

Across all experiments, reactions containing 5% blood consistently amplified faster than blood-free controls, regardless of the diluent used (Fig. 6B). This observation was unexpected given the presence of known amplification inhibitors in whole blood (Nwe et al., 2024). To further examine this effect, we compared amplification under three conditions: (i) template spiked into water, (ii) template spiked into 5% D-mannitol, and (iii) template spiked into 5% blood diluted in 5% D-mannitol. Fig. 6C shows standard calibration curves for all three conditions. Bland–Altman analysis showed a minimal difference in Tq between reactions prepared in water and 5% D-mannitol (1.1 min), indicating that D-mannitol alone did not substantially alter amplification kinetics relative to water (Fig. 6D). In contrast, reactions containing 5% blood diluted in 5% D-mannitol amplified, on average, 14.3 min faster than reactions containing 5% D-mannitol without blood (Fig. 6E). Together, these results demonstrate that the presence of diluted whole blood in 5% D-mannitol accelerates colorimetric LAMP amplification. The mechanism underlying this acceleration remains unclear and warrants further investigation in the future. It is possible that the anticoagulant present in blood collection tubes contributes to the observed acceleration, although this possibility was not examined in the present study.

## Discussion

In this study, we developed a colorimetric µPAD-LAMP assay targeting the ASFV VP72 gene and evaluated its suitability for PON deployment. The µPAD-LAMP assay achieved an LOD of 25 copies per reaction (3.3 copies/µL in the diluted sample) using synthetic DNA spiked into 5% diluted whole blood and was close to both the WOAH-recommended King’s qPCR assay (King et al., 2003) and the NAHLN qPCR assay (Zsak et al., 2005) under comparable conditions (Fig. 3, Fig. S4). Further, this analytical sensitivity is consistent with previously reported ASFV LAMP assays using synthetic targets (LOD: 1–50 copies/reaction; Table 1). However, unlike many previous reports (Table 1), the µPAD-LAMP sensor developed in this study requires minimal sample processing, consisting only of whole blood dilution prior to amplification. In addition, the assay enables both qualitative visual interpretation through color change and quantitative analysis using image processing. Therefore, the primary advancement of this work lies in simplifying sample preparation to enable direct LAMP detection from diluted whole blood while maintaining analytically relevant sensitivity. After accounting for dilution, the effective LOD corresponds to ∼67 copies/µL in whole blood, which is equivalent to ASFV blood viral loads observed at approximately 2–3 days post-infection in experimentally inoculated pigs (Lee et al., 2021).

In addition, the assay demonstrated high analytical specificity, with no cross-reactivity observed against nine common swine viral and bacterial pathogens (Fig. 4). High analytical specificity is particularly important for ASFV diagnostics because acute ASFV infection progresses rapidly (within few days) and its clinical and post-mortem manifestations cannot be reliably distinguished from other febrile hemorrhagic syndromes or bacterial septicemias of pigs by clinical or post-mortem examination alone (Lim et al., 2023; WOAH, 2022). Therefore, a highly specific molecular assay is essential to support accurate laboratory confirmation of ASFV and enable timely implementation of regulatory control measures, including movement restrictions and regional depopulation when necessary.

The real-time colorimetric tracking enables extraction of quantitative information from the colorimetric LAMP assay (Fig. 2D). However, compared to the NAHLN and King’s qPCR assays, the LOQ of 250 copies/reaction obtained with the paper LAMP assay was approximately ten times worse in the 5% blood matrix. Although quantitative thresholds such as LOQ are not typically required for ASF diagnosis, which primarily relies on qualitative detection due to stamping-out control policies, viral load measurements can provide additional value in specific contexts. In particular, quantitative data can support estimation of infection stage and improve epidemiological investigations, including inference of infection timing and transmission dynamics (Dixon et al., 2020; Fernández-Pinero et al., 2013; Lee et al., 2021). Thus, while less sensitive for quantification than qPCR, the ability to extract semi-quantitative information from a field-deployable paper LAMP assay may still offer practical benefits.

To evaluate operational transferability and suitability for PON deployment, we assessed assay performance in an external laboratory using operators unfamiliar with the µPAD-LAMP format but provided with specific written instruction manual (see SI, “ASF_Field_Manual.docx”) and a single on-site demonstration and training visit. Under these conditions, the assay achieved 90% analytical sensitivity and 92% specificity, representing an ∼8-10% reduction compared with the 100% sensitivity and analytical specificity obtained during expert-operated testing in our laboratory (Fig. 1). This decrease likely reflects workflow-dependent factors, including user handling variability (e.g., pipetting errors) and partial sample or reagent degradation during storage and transport, which become apparent when assays are transferred outside the developer laboratory. Importantly, this evaluation provided actionable insight into aspects of the workflow that require further standardization for PON use. Because ASFV is a foreign animal disease subject to strict regulatory and biosafety controls in the United States, evaluation using naturally infected clinical specimens was not feasible; therefore, assay performance was assessed using controlled, plasmid-spiked samples designed to approximate clinically relevant viral loads.

Further, we observed a counterintuitive matrix effect during assay optimization. Specifically, LAMP amplification proceeded faster in reactions containing whole blood diluted in 5% D-mannitol than in reactions containing water or D-mannitol alone (Fig. 6C). To our knowledge, there are no peer-reviewed reports describing blood-driven acceleration of LAMP amplification; instead, existing studies predominantly emphasize the inhibitory effects of blood components, such as hemoglobin and immunoglobulins, on nucleic acid amplification (Nwe et al., 2024). This observation contrasts with prevailing assumptions that whole blood acts as a strictly inhibitory matrix, particularly when combined with isotonic solutions, and warrants further investigation in the future to determine whether this effect is reproducible across blood sources, collection methods, and assay formats.

Future work will focus on further improving the diagnostic workflow by integrating sample collection, dilution, and loading into a single microfluidic chip coupled directly to the µPADs, thereby reducing user handling and operational variability. In parallel, several of these efforts are already underway, including the development of a circular microfluidic distribution chip and a water-free, portable heater–imager platform based on indium tin oxide (ITO) heating elements, designed to enable controlled, uniform heating without the need for a water bath (Raut et al., 2026b). Beyond workflow integration and hardware development, continued validation at endemic sites using naturally infected clinical specimens will be required to establish clinically relevant diagnostic performance, along with systematic evaluation of assay lifetime and reagent stability under realistic storage and environmental conditions.

## Conclusions

In this study, we developed a colorimetric µPAD-LAMP diagnostic platform for ASFV detection directly from minimally processed whole blood and evaluated its suitability for field deployment. The µPAD-LAMP system demonstrates three key advantages: (i) extraction-free detection of ASFV VP72 targets in 5% diluted whole blood with LOD of 25 copies/reaction (effective ∼67 copies/µL in whole blood), corresponding to early-stage infection (∼2–3 days post-infection); (ii) high specificity with no cross-reactivity against nine common swine viral and bacterial pathogens; and (iii) operational transferability, maintaining >90% analytical sensitivity and analytical specificity when deployed outside the developer laboratory.

The current system has two principal limitations: (i) diagnostic performance was evaluated using plasmid-spiked blood rather than naturally infected clinical specimens due to regulatory constraints associated with ASFV in the United States; and (ii) the sample handling steps introduce user-dependent variability that impacts performance when the assay is transferred beyond expert operators.

Future work will focus on two primary directions: (i) further integration of sample collection, dilution, and loading into a closed, microfluidic-assisted workflow coupled directly to the µPADs, alongside continued development of a water-free, portable heater–imager platform based on ITO or printed circuit board (PCB) heating elements to reduce contamination risk and improve usability; and (ii) expanded validation at endemic sites using naturally infected samples, including systematic evaluation of assay lifetime, reagent stability under realistic environmental conditions, and multi-user testing to establish clinically relevant diagnostic performance. Collectively, this work positions the µPAD-LAMP platform as a promising foundation for scalable, extraction-free molecular diagnostics for ASFV and other high-consequence veterinary pathogens.

## CRediT authorship contribution statement

**Bibek Raut:** Conceptualization, software, formal analysis, investigation, methodology, validation, visualization, writing – original draft preparation. **Gopal Palla:** Conceptualization, data curation, formal analysis, investigation, methodology, validation, writing – review and editing. **Nafisa Rafiq:** Data curation, formal analysis, investigation, methodology, validation, writing – review and editing. **Jiangshan Wang:** Data curation, formal analysis, investigation, methodology, writing – review and editing. **Virendra Kumar:** Conceptualization, Data curation, methodology, writing – review and editing. **Mohamed S. Kamel:** Investigation, Data curation, methodology. **Dan Van Nguyen:** Data curation, methodology. **Saraswathi Lanka:** Data curation, investigation, validation, writing – review and editing. **Carol W. Maddox:** Supervision, writing – review and editing**. Darryl Ragland**: Supervision, funding acquisition. **J. Alex Pasternak:** Conceptualization, Supervision. funding acquisition, validation, writing – review and editing. **Mohit S. Verma:** Conceptualization, supervision, project administration, funding acquisition, validation, and writing – review & editing. All authors have reviewed the manuscript and approved the final version.

## Declaration of generative AI and AI-assisted technologies in the writing process

During the preparation of this work, the authors used ChatGPT to check for grammar errors and improve their academic writing language as well as debugging software. After using this tool/service, the authors reviewed and edited the content as needed and take full responsibility for the content of the publication.

## Use of animal in the experiment

All animal procedures were approved by the Purdue University Institutional Animal Care and Use Committee (IACUC number: IPROTO2205002265).

## Declaration of competing interest

The authors declare the following financial interests/personal relationships which may be considered as potential competing interests:

Mohit S. Verma reports a relationship with Krishi, Inc. that includes: board membership, equity or stocks, and funding grants. Krishi, Inc. did not fund this work. Mohit S. Verma also has a relationship with Simply Experiment LLC, that includes: equity and ownership. Simply Experiment LLC did not fund this work. The other authors declare that they have no known competing financial interests or personal relationships that could have appeared to influence the work reported in this paper.

## Acknowledgements

We are grateful to Alyssa A. Smith, Dana Jeon, and Kaylyn Rudy for their help with collecting the blood samples.

## Funding

This project is supported by United States Department of Agriculture Animal and Plant Health Inspection Service (APHIS) through the National Animal Health Laboratory Network (Sponsor Award # AP22VSD&B000C022). The findings and conclusions in this publication are those of the authors and should not be construed to represent any official USDA or U.S. Government determination or policy. The project is also partially supported by Purdue University’s 2025 Bridge Funding program.

## Appendix A. Supplementary data

The following are the Supplementary data to this article: Supporting information document ASF_Field_Manual.docx VP72_TOPII_Template_Sequence.xlsx

## Data availability

This article includes Supporting Documents and a Mendeley dataset. A temporary access link to Mendeley data is provided below for peer review purposes; the final public link will be made available upon manuscript acceptance: https://data.mendeley.com/preview/fwg5yjpygg?a=6769860b-79d2-4174-b248-f51124b1b4c4

## Notes

https://data.mendeley.com/preview/fwg5yjpygg?a=6769860b-79d2-4174-b248-f51124b1b4c4

