## Supporting Information for "A rapid, field-deployable paper-based biosensor for the detection of African swine fever virus in whole blood"

for

### Note 1.1: ASFV synthetic plasmid targets synthesis and quantification

Synthetic DNA plasmid targets corresponding to the ASFV B646L (VP72) and TOPII genes were designed based on the NCBI reference sequences MG209614.1 (B646L) and Z14245.1 (TOPII). A 1,953-bp fragment of the B646L gene was flanked by EcoRI and KpnI restriction sites and cloned into the pUC57 vector, generating a 4,651-bp synthetic plasmid. Similarly, a 3,588-bp fragment of the TOPII gene was flanked by EcoRV and BamHI restriction sites and cloned into pUC57 to generate a 6,280-bp synthetic plasmid. Both plasmids were synthesized and sequence-verified by GenScript Biotech Corporation and used as standardized positive-control templates for LAMP assay development. The full sequences are provided in the Supporting Information (“VP72_TOPII_Template_Sequence.xlsx”). Digital PCR (dPCR) was performed to quantify the synthetic templates using previously reported primers and probes for VP72 (King et al., 2003) and a custom-designed primer set for TOPII using similar protocol to reported earlier (Ahmed et al., 2025a; Raut et al., 2026a). Briefly, each dPCR assay targeting the synthetic DNA standard was assembled in a final reaction volume of 40 μL. The mixture contained 10 μL of 4× Probe PCR Master Mix (Qiagen, 250102; yielding a final 1× concentration), 4 μL of a 10× primer–probe mix (final concentrations: 0.8 μM each forward and reverse primer, and 0.4 μM FAM-labeled probe; Table S1), 0.5 μL of EcoRI-HF restriction enzyme (New England Biolabs, R3101S), 20.5 μL of nuclease-free water, and 5 μL of DNA template. Prepared reactions were loaded onto a 26K 24-well Nanoplate (Qiagen, 250001) and analyzed using the QIAcuity One 5-plex digital PCR system (Qiagen, 911021). Thermal cycling conditions included an initial denaturation step at 95 °C for 2 min, followed by 40 amplification cycles consisting of 95 °C for 15 s, 55 °C for 15 s, and 60 °C for 30 s. Absolute quantification of DNA copy numbers was performed using the QIAcuity Software Suite (Qiagen). Following quantification, the synthetic DNA stock was stored at −80 °C until needed. dPCR Primer and probe sequences are listed in Table S1.

### Note 1.2: ASFV LAMP primer design, synthesis, and screening

Six sets of LAMP primers were designed for each target. One previously published LAMP primer set targeting *B646L* (VP72) (Y. Wang et al., 2021) and one targeting *TOPII* (James et al., 2010) were included as reference controls. LAMP primers were designed for each gene using PrimerExplorer V5 (<https://primerexplorer.jp/lampv5e/index.html>) and ordered from Integrated DNA Technologies, Inc., USA. Information for all 14 primer sets is provided in Table S2, while the final primer sets used in paper LAMP experiments are listed in Table 2. Primer sets were named using the following convention: in the name ASFV.VP72.1, “ASFV” denotes the target organism, “VP72” indicates the target gene/protein, and “1” represents the primer set number.

Fluorescent LAMP reactions were performed using the primer sets listed in Table S2. Each reaction mixture contained 1× LAMP primer mix (1.6 μM FIP and BIP, 0.2 μM F3 and B3, and 0.4 μM LF and LB), 2.5 µl of WarmStart® 2× LAMP Master Mix (E1700; New England Biolabs, USA), and 2.5 µl of a 2× fluorescent dye solution prepared by diluting the supplied 50× LAMP fluorescent dye with nuclease-free water. Reaction mixtures (5 µL) were dispensed into wells of a 384-well plate (Fisher Scientific, 44-832-73, USA). Plasmid DNA was then added at defined copy numbers using an Echo acoustic liquid handler (Beckman Coulter, Echo 650, USA) in 2.5-nL increments. Target copy numbers were achieved by transferring nanoliter volumes from two plasmid stocks (4 copies/nL and 40 copies/nL), with the lower-concentration stock used for low copy inputs (10, 50, 100, and 500 copies/reaction) and the higher-concentration stock used for high copy inputs (1000, 5000, and 10000 copies/reaction). No-template control (NTC) reactions did not receive an equivalent volume of template diluent because the template addition volumes (2.5–250 nL) are negligible relative to the total reaction volume. The plate was sealed using a PlateLoc Thermal Microplate Sealer (Agilent, PlateLoc, USA) and incubated in a qTower 384 real-time PCR system (Analytic Jena, Germany) at 65 °C for 60 min with a temperature ramp rate of 0.1 °C s⁻¹. Fluorescence signals were monitored in real time using the FAM detection channel, with data acquisition performed at 60-s intervals throughout the incubation period.

The LAMP primers were sorted based on five metrics using the Primer Scoring software (https://github.itap.purdue.edu/VermaLab/PrimerScoring) (Kamel et al., 2025; Pascual-Garrigos et al., 2021). The software scores primer sets using five quantitative performance parameters derived from replicate amplification data: (i) average maximum fluorescence intensity, (ii) standard deviation of maximum intensity, (iii) average reaction time, (iv) standard deviation of reaction time, and (v) false positives. Positive reactions are first identified algorithmically from the fluorescence time series based on signal shape and intensity thresholds, and reaction time is defined as the point of maximum second derivative. For each primer set, these metrics are calculated across replicates, with intensity and speed rewarded and variability penalized. Each parameter is normalized relative to the range observed across all primer sets and weighted according to predefined importance. False positives are additionally penalized in a rank-based manner across replicates, with earlier and stronger false amplification incurring larger penalties. The weighted contributions are summed to produce an overall score, and primer sets that do not achieve amplification in all replicates are assigned a score of zero (Kamel et al., 2025).

### Note 1.3: µPAD and acrylic cartridge fabrication

Microfluidic paper-based analytical devices (µPAD) (3 × 3 mm) were fabricated following a previously reported protocol (Ahmed et al., 2025a; Raut et al., 2026a). Briefly, chromatographic grade 222 paper (Ahlstrom-Munksjö, Finland) and polystyrene spacer sheets (HIPS Litho Grade, Tekra, USA) were cut into 3 mm–wide strips using a leather strip-cutting machine (Zhixumm, Amazon, USA) with the blade spacing set to 3 mm. A transparent polyester film (MELINEX® 454, Tekra, USA) was taped with double-sided adhesive (ARclean® 90178, USA), after which paper and spacer strips were sequentially assembled onto the adhesive surface. The assembled spacer–paper structure was then aligned and cut using the leather cutter into two-, three-, or four-pad strips and separated using scissors.

Acrylic cartridges for housing the µPAD strips were fabricated following a previously reported design (Raut et al., 2026b, 2026a). Cartridge designs were created using SolidWorks, exported as DXF files, and edited in Adobe Illustrator (Adobe Inc., USA) prior to conversion to SVG format for laser cutting. Cartridges were cut from 1.5-mm-thick acrylic sheets (Outus, Amazon, USA) using a 40 W Glowforge Plus laser cutter (Glowforge Inc., USA) with optimized cutting parameters. After fabrication, acrylic components were rinsed with reverse osmosis (RO) water, treated sequentially with 70% ethanol and RNase AWAY™ (Thermo Fisher Scientific, USA), and wiped with lint-free Kimwipes® (Kimberly-Clark, USA). The cartridge design was readily adapted to accommodate different µPAD strip formats (2, 3, or 4-pad configurations) and slot geometries (full or half).

### Note 1.4: Preparation of colorimetric paper LAMP and assembly of µPADs into cartridge

The colorimetric LAMP formulation for the paper-based assay was adapted from a previously reported protocol used for detection of Severe Acute Respiratory Syndrome Coronavirus 2 (SARS-CoV-2) (Davidson et al., 2021; J. Wang et al., 2021). A homemade 2× LAMP master mix was prepared using KCl (Sigma-Aldrich, USA), MgSO₄ (Sigma-Aldrich, USA), a dNTP mixture (Fisher Scientific, USA), dUTP (Fisher Scientific, USA), Antarctic Thermolabile (Ant) UDG (New England Biolabs, USA), Bst 2.0 DNA polymerase (New England Biolabs, USA), phenol red as a pH-sensitive indicator (Sigma-Aldrich, USA), Tween-20 (Sigma-Aldrich, USA), and nuclease-free water (Fisher Scientific, USA). The identity and concentrations of all components used in the 2× LAMP mix are provided in Table S3.

The final 200 µL LAMP reaction mixture was prepared by combining the 125 μL of 2× LAMP master mix with target-specific LAMP primer sets (25 μL 10× primer mix (16 μM FIP/BIP, 2 μM F3/B3, and 4 μM LF/LB; final concentrations 1.6 μM FIP/BIP, 0.2 μM F3/B3, 0.4 μM LF/LB), additional 0.67 μL Bst 2.0 DNA polymerase, 1µL betaine (Sigma-Aldrich, USA), 3.13 μL bovine serum albumin (BSA; Sigma-Aldrich, USA) (40 mg/mL; Sigma-Aldrich, A2153), 36.0 μL trehalose (1.75 M; Thermo Scientific Chemicals, 182550250), and 9.2 μL nuclease-free water. Final reaction component concentrations are summarized in Table S4.

Aliquots (7.5 µL) of the final LAMP master mix were pipetted onto individual µPADs and allowed to dry inside a PCR workstation under ambient conditions for approximately 2 h. Following drying, the reagent-loaded µPAD strips were carefully transferred using sterile tweezers and assembled into the acrylic cartridges as described above. One side of each cartridge was sealed with PCR plate sealing film (Fisher Scientific, USA), to which the µPAD strips were attached. The assembled cartridges containing µPAD strips were used either immediately or stored at −20 °C for up to 1 week.

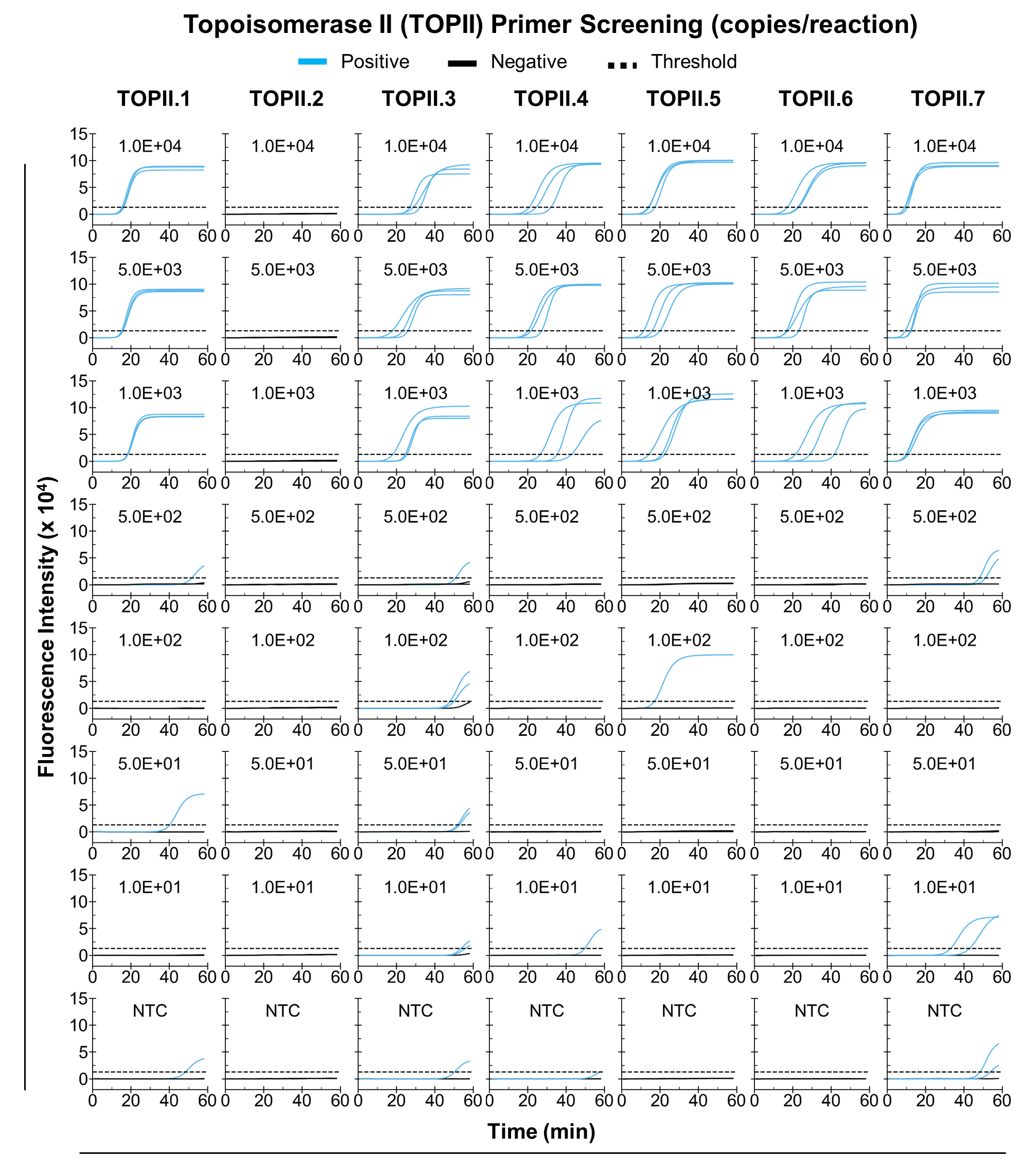

Figure S1. Primer screening using synthetic Topoisomerase II (TOPII) DNA targets showing positive and negative amplification results based on a fixed positivity threshold. NTC, no-template control.

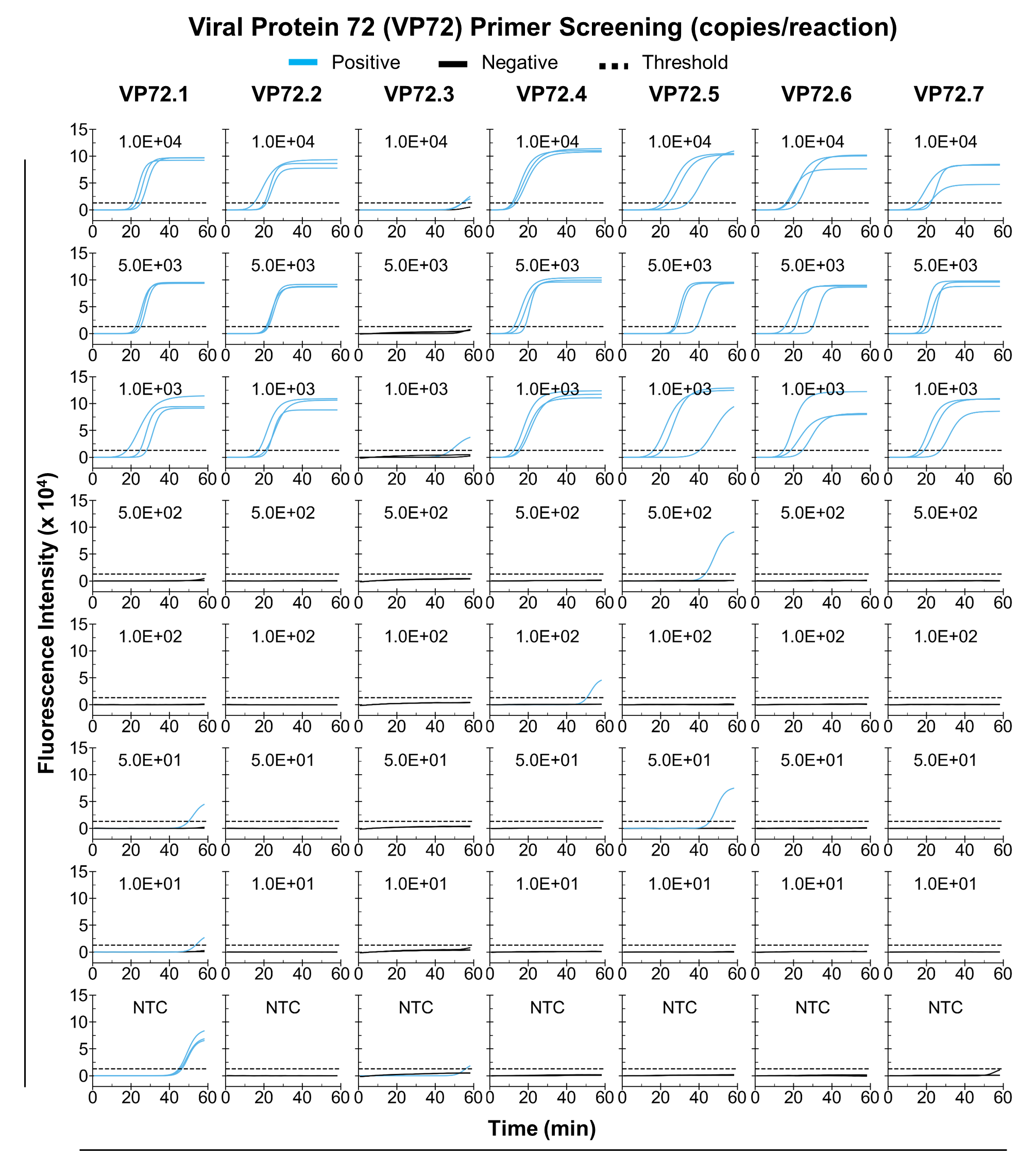

Figure S2. Primer screening using synthetic Viral Protein 72 (VP72) DNA targets showing positive and negative amplification results based on a fixed positivity threshold. NTC, no-template control.

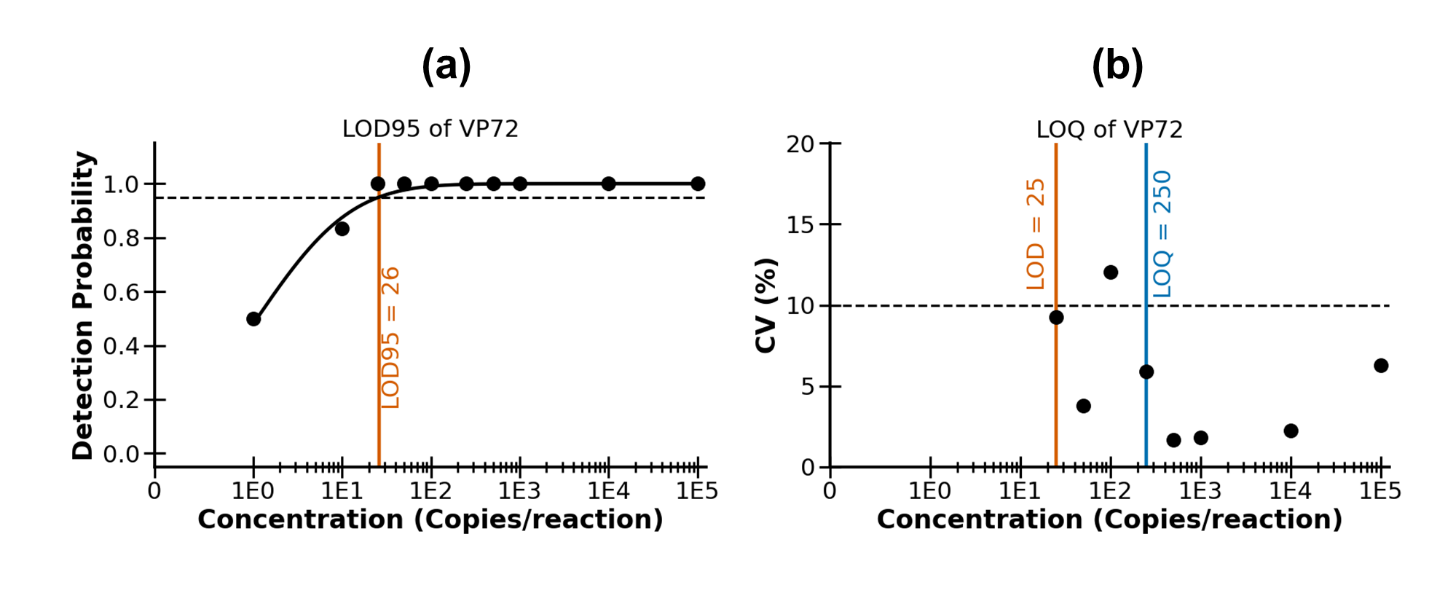

Figure S3. Limit of detection (LOD) and limit of quantification (LOQ) determination for the VP72 target. (a) Probit fit showing the 95% detection probability from serially diluted standards, with LOD95 defined as the lowest concentration that crosses the 95% detection threshold. (b) A CV cutoff of 10% was used to define the LOQ as the lowest concentration that maintained a CV within this threshold. The LOD shown here represents the final LOD determined after further evaluation of the preliminary LOD95 at 25 copies per reaction, where more than 19 of 20 technical replicates were positive and no negative controls showed amplification.

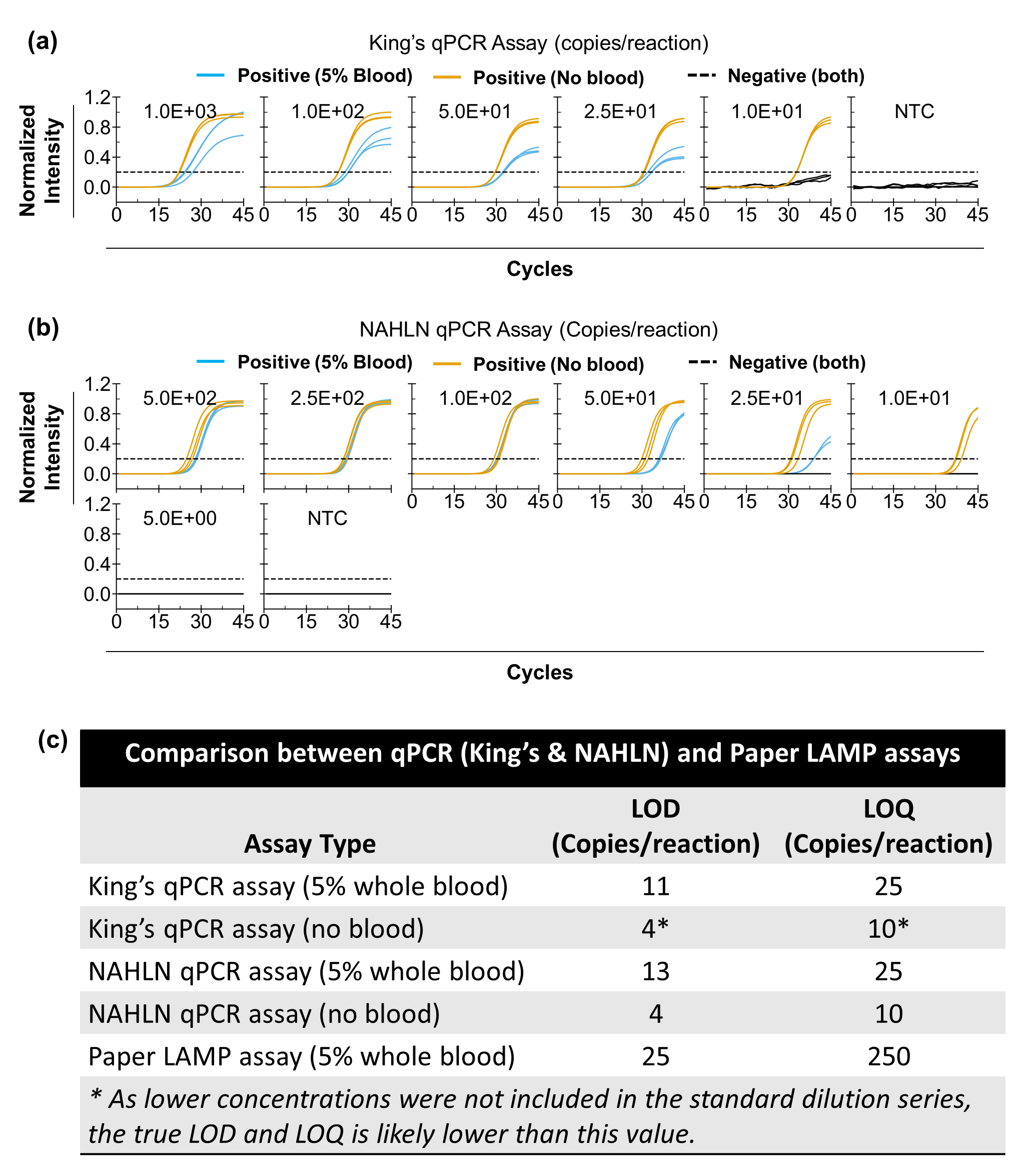

Figure S4. Analytical sensitivity evaluation of the NAHLN and King’s qPCR assays using VP72 targets. VP72 plasmid DNA was spiked at concentrations ranging from 1,000 copies/reaction to 0 (no-template control (NTC)) into either whole blood diluted to 5% (v/v) in 5% D-mannitol (corresponding to a final blood concentration of 1% (v/v) in the qPCR reaction) or 5% D-mannitol alone. (a) Serial diluted standard concentrations and their corresponding normalized fluorescence intensity for the King’s qPCR assay and for (b) NAHLN qPCR assay. (c) Comparison of the limit of detection (LOD) at 95% detection probability and limit of quantification (LOQ) across the assays and the paper LAMP assay.

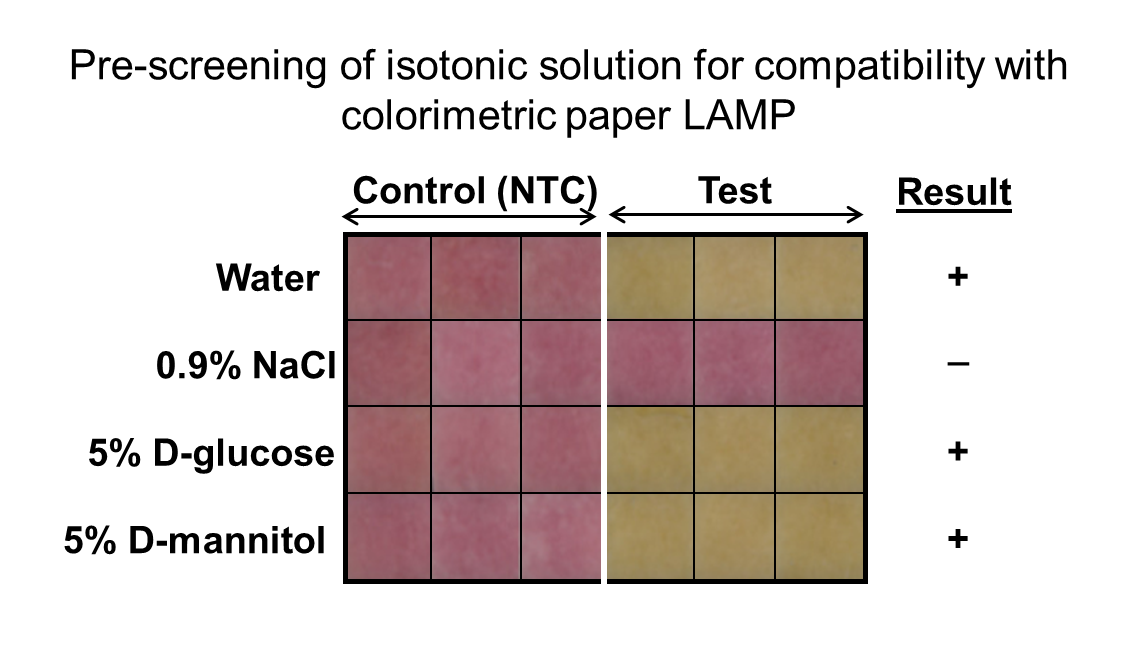

Figure S5. End result (t = 60 min) of paper LAMP when tested with sample spiked in either water, 0.9% NaCl, 5% D-glucose, or 5% D-mannitol.

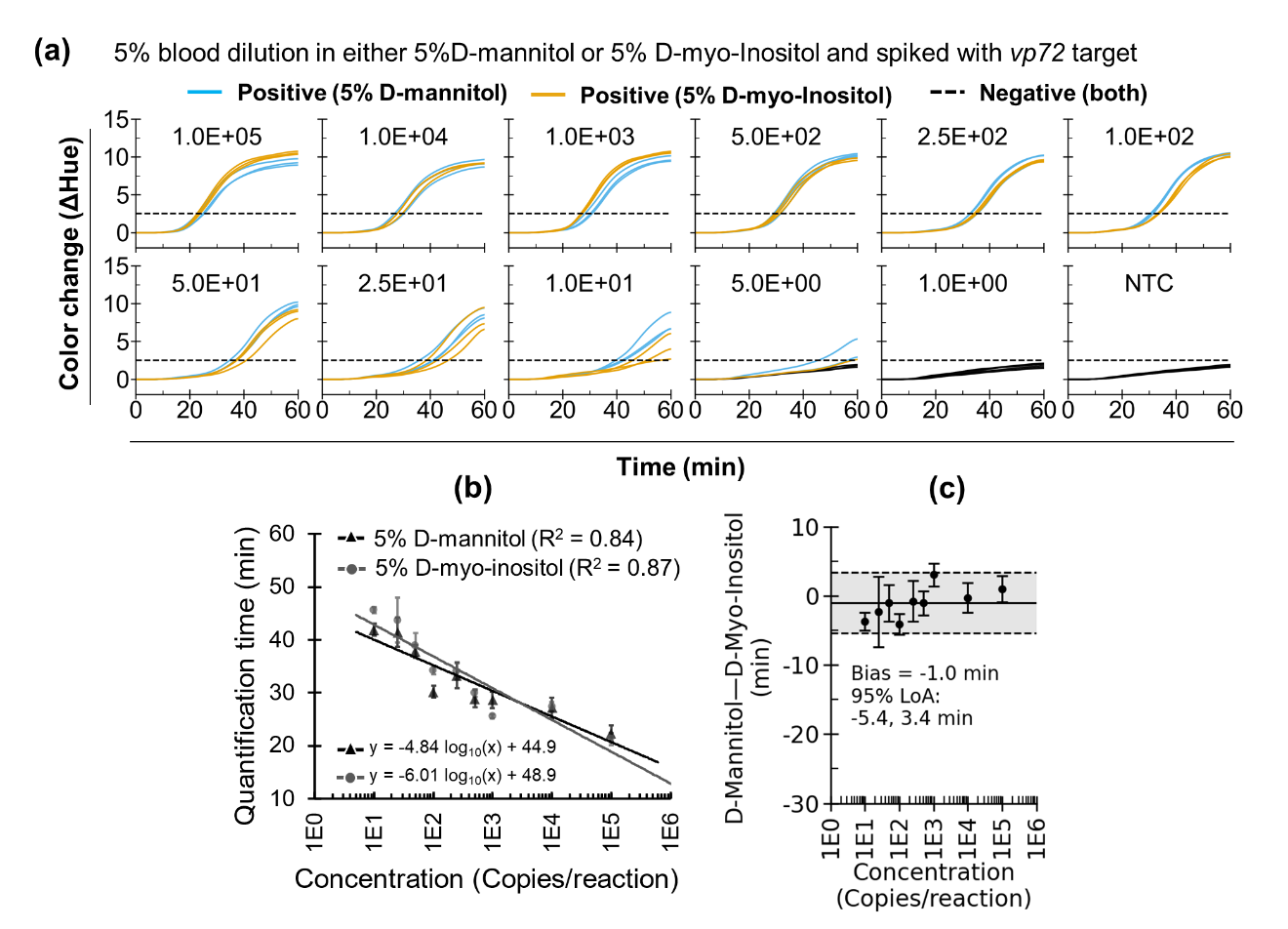

Figure S6. Comparison of 5% D-myo-inositol and 5% D-mannitol for blood dilution. (a) Net hue–versus–time plots for serially diluted concentrations of synthetic VP72 DNA spiked into 5% blood diluted with either 5% D-myo-inositol or 5% D-mannitol, along with no-template controls (NTCs). Freshly collected blood (<24 h) was used for all reactions. (b) Quantification time (Tq) versus concentration derived from (a) for the two dilution conditions, 5% D-myo-inositol and 5% D-mannitol. (c) Bland–Altman plot showing the Tq difference of 5% D-mannitol minus 5% D-myo-inositol as a function of concentration.

Table S1: dPCR/qPCR primer sets for detection of various swine pathogens.

| **Target** | **Forward primer, Reverse primer, and Probe (5' to 3')** | | **References** |
| --- | --- | --- | --- |
| ASFV_VP72  (Set 1) | Fwd | CTGCTCATGGTATCAATCTTATCGA | (King et al., 2003) |
|  | Rev | GATACCACAAGATC(AG)GCCGT |  |
|  | Probe | /56-FAM/CCACGGGAGGAATACCAACCCAGTG/3IABkFQ/ |  |
| ASFV_VP72  (Set 2) | Fwd | C**T**TCGGCGAGCGCTTTATCAC | With slight modification shown in bold blue (Zsak et al., 2005) |
|  | Rev | GGAAA**T**TCATTCACCAAATCCTT |  |
|  | Probe | /56-FAM/ CGATGCAAGCTTTAT/3IABkFQ/ |  |
| ASFV_TOPII | Fwd | TCACAGTGGGCCATTCTTAC | This study |
|  | Rev | CATTTCCCTGGAGGGTGTAATC |  |
|  | Probe | /56-FAM/CAGCGCCTGGATATCATTGAGCCT/3IABkFQ/ |  |
| *Bordetella bronchiseptica* | Fwd | ACTATACGTCGGGAAATCTGTTTG | (Fastrès et al., 2020) |
|  | Rev | CGTTGTCGGCTTTCGTCTG |  |
|  | Probe | /56-FAM/CGGGCCGATAGTCAGGGCGTAG/3IABkFQ/ |  |
| *Pasteurella multocida* | Fwd | GGAAGCCTTCCAAGCAGAATTTG | (Tocqueville et al., 2017) |
|  | Rev | CCGCAATAGCTTTACCCATTACAG |  |
|  | Probe | /56-FAM/CGGCAGCAACCCGTTTCGGTTCAG/3IABkFQ/ |  |
| *Actinobacillus*  *pleuropneumoniae* | Fwd | GGGGACGTAACTCGGTGATT | (Tobias et al., 2012) |
|  | Rev | GCTCACCAACGTTTGCTCAT |  |
|  | Probe | /56-FAM/CGGTGCGGACACCTATATCT/3IABkFQ/ |  |
| *Streptococcus suis* | Fwd | GTCATGGACTCGTGAAGAAGTAG | (Yi et al., 2022) |
|  | Rev | GTAGTCTGTACCAAGGTCGTATTT |  |
|  | Probe | /56-FAM/TTGGCTGTGTTGAAGATGTTGGCC/3IABkFQ/ |  |
| *Actinobacillus suis* | Fwd | GAGCTGGGAAGCTCGACTAT | (Kariyawasam et al., 2011) |
|  | Rev | CCCCCATCTTCAAACAGGAT |  |
|  | Probe | /56-FAM/AGCTAACGACAAGTAGGGCG/3IABkFQ/ |  |
| Porcine circovirus type 2d (PCV2d) | Fwd | CCCTGTCACCCTGGGTGAT | (Li et al., 2018) |
|  | Rev | CCTGTGCCCTTTGAATACTACAGA |  |
|  | Probe | /56-FAM/TAAGGTTGAATTCTGGCCCTGCTCCC/3IABkFQ/ |  |
| Porcine Respiratory and Reproductive Syndrome Virus 2 (PRRSV2, strain: VR-2332 ) | Fwd | CCGCGTAGAACTGTGACAAC | (Wernike et al., 2012) |
|  | Rev | TCCAGGATGCCCATGTTCTG |  |
|  | Probe | /56-FAM/ACGCACCAGGATGAGCCTCTGGAT/3IABkFQ/ |  |
| Porcine Respiratory Coronavirus (PRCoV) | Fwd | ATAGACAAACTCGCTATCGC | (Keep et al., 2022) |
|  | Rev | CAACCCAGACAACTCCATC |  |
|  | Probe | /56-FAM/TAGGCACTGGACCTCATGCA/3IABkFQ/ |  |
| Influenza A (H1N1) - Swine origin | Fwd | TGTGAATCACTCTCCACAGCAAGC | (Wang et al., 2009) |
|  | Rev | ATTGGGCCATGAACTTGTCTTGGG |  |
|  | Probe | /56-FAM/CAGACAATGGAACGTGTTACCCAGGA/3IABkFQ/ |  |

Table S2: LAMP primer sets designed for detection of *B646L* (VP72) and *TOPII* genes.

| **Primers** | **sequence (5' to 3')** | **Reference** |
| --- | --- | --- |
| ASFV_TOPII.1_F3 | GGCGCAAAATTTTAGCCGG | (James et al., 2010) |
| ASFV_TOPII.1_B3 | GCCGAAGCTTCCTATGCC |  |
| ASFV_TOPII.1_FIP | GCAACGTAGCCCCCGAACTGGAAATGCTTCGCYTCCAACA |  |
| ASFV_TOPII.1_BIP | ATCACCATGGCGACATGTCGTGGATAGAGGTGGGAGGAGC |  |
| ASFV_TOPII.1_LF | AAAAACCTTTCGTTCACGGT |  |
| ASFV_TOPII.1_LB | AAAAGCCGCCCAGTATTACC |  |
| ASFV_TOPII.2_F3 | GTGACCGACAAATTCCTTGC | (This study) |
| ASFV_TOPII.2_B3 | ACCTTTCGTTCACGGTTGT |  |
| ASFV_TOPII.2_FIP | AAGTTGGGTATTTGCCGCTCGACCTGCACCTCCAGGTAGA |  |
| ASFV_TOPII.2_BIP | TTAGACGGGATGACCCGGGCTGGAGGCGAAGCATTTCAAC |  |
| ASFV_TOPII.2_LF | AGCTTGTAGGCCTTGGTG |  |
| ASFV_TOPII.2_LB | GCGGCGCAAAATTTTAGCC |  |
| ASFV_TOPII.3_F3 | ACACCAAGGCCTACAAGCT | (This study) |
| ASFV_TOPII.3_B3 | CGACATATCGCCATGGTGAT |  |
| ASFV_TOPII.3_FIP | CCGGCTAAAATTTTGCGCCGCCGAGCGGCAAATACCCAAC |  |
| ASFV_TOPII.3_BIP | CTTCGCCTCCAACAACCGTGAAACATGTGATCCGCAACGT |  |
| ASFV_TOPII.3_LF | CGGGTCATCCCGTCTAAGAA |  |
| ASFV_TOPII.3_LB | ACGAAAGGTTTTTCAGTTTGGG |  |
| ASFV_TOPII.4_F3 | CTACTGCACTACGCAATAG | (This study) |
| ASFV_TOPII.4_B3 | CCGAACTACTGTAGTCAATG |  |
| ASFV_TOPII.4_FIP | GCCGAATCGTTTCAAGTGACCGTAAGATCACTGTACTTCC |  |
| ASFV_TOPII.4_BIP | GCACTTACGTCGTCTCAGAGGTAAGCAACCGTAGGAAC |  |
| ASFV_TOPII.4_LF | TGTAATTAGAAGGCCGTAGC |  |
| ASFV_TOPII.4_LB | ATTACTATTACGGAGCTTCCTC |  |
| ASFV_TOPII.5_F3 | TATTACAGCTACGGCACT | (This study) |
| ASFV_TOPII.5_B3 | TCTTGTTCTTCAGTCTCCT |  |
| ASFV_TOPII.5_FIP | ATGCCATGCGGTTACTCGTTACGGAGCTTCCTCTAC |  |
| ASFV_TOPII.5_BIP | ACAGTAGTTCGGAAACCATTGAACGATACGGTTGAGACTAT |  |
| ASFV_TOPII.5_LF | TAAGCAACCGTAGGAACAC |  |
| ASFV_TOPII.5_LB | TTCTGGTGAAGTTAAAGCCA |  |
| ASFV_TOPII.6_F3 | CCAATACTATTACAGCTACGG | (This study) |
| ASFV_TOPII.6_B3 | TCTTGTTCTTCAGTCTCCT |  |
| ASFV_TOPII.6_FIP | CGATGTAAGCAACCGTAGGAAACTTACGTCGTCTCAGAG |  |
| ASFV_TOPII.6_BIP | ATCGAGTAACCGCATGGCCTTCACCAGAATTTCAATGG |  |
| ASFV_TOPII.6_LF | GAGGAAGCTCCGTAATAGTAAT |  |
| ASFV_TOPII.6_LB | TCATTGACTACAGTAGTTCGG |  |
| ASFV_TOPII.7_F3 | CGCGCAAAAGTATGTTCAAC | (This study) |
| ASFV_TOPII.7_B3 | TCCAGGCGGAAAGATCTG |  |
| ASFV_TOPII.7_FIP | GATACGTGTAAACATTTCCCTGGTTTTTAACCAGCGCCTGGATATC |  |
| ASFV_TOPII.7_BIP | CTCATGCCCGTATACCAGGTTTTTCTTGCTCCGTTTCAGACAG |  |
| ASFV_TOPII.7_LF | GTGTAATCGTAGGAGGCTCAA |  |
| ASFV_TOPII.7_LB | AACTAGGGTATGCGCAGC |  |
| **Primers** | **sequence (5' to 3')** | **Reference** |
| ASFV_VP72.1_F3 | ACGGACGCAACGTATCTGGA | (Y. Wang et al., 2021) |
| ASFV_VP72.1_B3 | TTGCGTCTACTGGGGCG |  |
| ASFV_VP72.1_FIP | CTTAATCCAGAGCGCAAGAGGGGTTTTTCTGGACATAAGACGTAATGTTCAT |  |
| ASFV_VP72.1_BIP | CCCTTCGGCGAGCGCTTTATTTTTGGAAATTCATTCACCAAATCCTTT |  |
| ASFV_VP72.1_LF | GGCTGATAGTATTTAGGGGTTTGA |  |
| ASFV_VP72.1_LB | TATCACCATAAAGCTTGCATCGC |  |
| ASFV_VP72.2_F3 | TCCAGGAAATTCATTCACC | (This study) |
| ASFV_VP72.2_B3 | CGAGAACTCTCACAATATCC |  |
| ASFV_VP72.2_FIP | TCTTGCGCTCTGGATTAAGTTGGAATGGATACTGAGGGA |  |
| ASFV_VP72.2_BIP | GGTTTGAGGTCCATTACAGCTCAAGATATTACTCCTATCACGG |  |
| ASFV_VP72.2_LF | GAGAACGTGAACCTTGCTA |  |
| ASFV_VP72.2_LB | TATGTCCAGATACGTTGCG |  |
| ASFV_VP72.3_F3 | AGTAGTAAACCAAGTTTCGG | (This study) |
| ASFV_VP72.3_B3 | GCTTACCTTTGGTATTCCC |  |
| ASFV_VP72.3_FIP | TTCCTCGCAACGGATATGACAGGATCTACAAGCGTGTA |  |
| ASFV_VP72.3_BIP | ACGTTTGAAGCTGCCCATGTGCATGTCATTCATCCT |  |
| ASFV_VP72.3_LF | AACCAAACACCCTTAGAGG |  |
| ASFV_VP72.3_LB | CTGAATCGGAGCATCCTG |  |
| ASFV_VP72.4_F3 | GGTAATGTGATCGGATACG | (This study) |
| ASFV_VP72.4_B3 | CAACAATAACCACCACGA |  |
| ASFV_VP72.4_FIP | TCACCACGCAGAGATAAGCATAGATGAACATGCGTCTG |  |
| ASFV_VP72.4_BIP | ATGTCCGAACTTGTGCCAAACCTACCTGGAACATCTC |  |
| ASFV_VP72.4_LF | CAGGATAGAGATACAGCTCTTC |  |
| ASFV_VP72.4_LB | CTCGGTGTTGATGAGGATT |  |
| ASFV_VP72.5_F3 | AAGATCAGCCGTAGTGATA | (This study) |
| ASFV_VP72.5_B3 | CGTATCCGATCACATTACC |  |
| ASFV_VP72.5_FIP | GTGCGATGATGATTACCTTTGCCTCTGGATACGTTAATATGACC |  |
| ASFV_VP72.5_BIP | CCTCCGTAGTGGAAGGGTCTGCTCATGGTATCAATCTT |  |
| ASFV_VP72.5_LF | CACGGGAGGAATACCAAC |  |
| ASFV_VP72.5_LB | ATGTAAGAGCTGCAGAACTT |  |
| ASFV_VP72.6_F3 | ACGTTTTCATAAAGTCGTTCTCC | (This study) |
| ASFV_VP72.6_B3 | CTTCAAACGTTTCCTCGCAA |  |
| ASFV_VP72.6_FIP | TTTGGAAGACCCATTGTACCCTTTTTGGTATTCGCAGTAGTAAACCAAG |  |
| ASFV_VP72.6_BIP | GGATCTACAAGCGTGTAAACGTTTTTCGGATATGACTGGGACAACC |  |
| ASFV_VP72.6_LF | GGCACAAAGAATGCGTACC |  |
| ASFV_VP72.6_LB | CGCCCTCTAAGGGTGTTT |  |
| ASFV_VP72.7_F3 | GCGTTGTGACATCCGAACTATA | (This study) |
| ASFV_VP72.7_B3 | CAACCAAACACCCTTAGAGG |  |
| ASFV_VP72.7_FIP | AAACTTGGTTTACTACTGCGAATACTTTTTGTCTAGGGAATTTCCATTTACATCG |  |
| ASFV_VP72.7_BIP | GGTACGCATTCTTTGTGCCTTTTTCGTTTACACGCTTGTAGATCC |  |
| ASFV_VP72.7_LF | CCCGGAGAACGACTTTATGAAA |  |
| ASFV_VP72.7_LB | GGGTACAATGGGTCTTCCAAA |  |

Table S3. Reagents used in preparing homemade 2X master mix.

| **Homemade 2X Mix** | | | | | | |
| --- | --- | --- | --- | --- | --- | --- |
| Components | Volume | Unit | Stock concentration | Unit | Final concentration | Unit |
| KCL | 100 | µL | 1000 | mM | 100 | mM |
| MgSO4 | 160 | µL | 100 | mM | 16 | mM |
| dNTPs | 280 | µL | 10 | mM | 2.8 | mM |
| Bst2.0 | 5.4 | µL | 120 | U/µL | 0.648 | U/µL |
| Rtx | 40 | µL | 15 | U/µL | 0.6 | U/µL |
| Phenol red | 20 | µL | 25 | mM | 0.2 | mM |
| dUTPs | 2.8 | µL | 100 | mM | 0.28 | mM |
| Ant UDG | 0.4 | µL | 1 | U/µL | 0.0004 | U/µL |
| Tween | 100 | µL | 20 | % | 2 | % |
| Water | 291.4 | µL | - | - | - | - |
| Total |  |  |  |  | 1000 | µL |

Table S4. Reagents used in preparing final reaction mix on paper.

| **Final Reaction Mix on paper** | | | | | | |
| --- | --- | --- | --- | --- | --- | --- |
| Components | Volume | Unit | Stock concentration | Unit | Final concentration | Unit |
| 2X Mix | 125 | µL | - | - | - | - |
| Primer Mix | 25 | µL | - | - | - | - |
| Bst2.0 | 0.67 | µL | 120 | U/ µL |  |  |
| Betaine | 1 | µL | 5 | M | 20 | mM |
| BSA | 3.13 | µL | 40 | mg/mL | - | - |
| Trehalose | 36 | µL | 62 | %(w/v) | ~11 | %(w/v) |
| Water | 9.2 | µL | - | - | - | - |
| Total |  |  |  |  | 200 | µL |

| Table S5. TOPII Primer Ranking | | | | | | | | |
| --- | --- | --- | --- | --- | --- | --- | --- | --- |
| Primer  Set | **True**  **Positive** | **Fluorescence**  **Intensity** | | **Run time (min)** | | **False**  **Positives** | **Overall**  **Score**  **(0-100)** | **Ranking** |
|  |  | **Max** | **SD** | **Mean** | **SD** |  |  |  |
| TOPII.5 | 3 | 119,031 | 5,481 | 18.7 | 4.0 | 0 | 90.1 | 1 |
| TOPII.1 | 3 | 84,626 | 2,515 | 17.0 | 0.0 | 1 | 84.1 | 2 |
| TOPII.6 | 3 | 104,521 | 6,299 | 30.3 | 8.2 | 0 | 77.2 | 3 |
| TOPII.3 | 3 | 88,797 | 12,051 | 22.0 | 3.6 | 1 | 77.1 | 4 |
| TOPII.7 | 3 | 92,238 | 2,622 | 9.0 | 0.8 | 2 | 65.5 | 5 |
| TOPII.4 | 3 | 100,583 | 21,735 | 40.0 | 13.9 | 0 | 62.3 | 6 |
| TOPII.2 | 0 | 0 | 0 | 0.0 | 0.0 | 0 | 0 | 7 |

| Table S6. VP72 Primer Ranking | | | | | | | | |
| --- | --- | --- | --- | --- | --- | --- | --- | --- |
| Primer  Set | **True**  **Positive** | **Fluorescence**  **Intensity** | | **Run time (min)** | | **False**  **Positives** | **Overall**  **Score**  **(0-100)** | **Ranking** |
|  |  | **Max** | **SD** | **Mean** | **SD** |  |  |  |
| VP72.4 | 3 | 117,161 | 6,521 | 13.7 | 1.2 | 0 | 100.0 | 1 |
| VP72.2 | 3 | 101,035 | 11,312 | 19.3 | 1.7 | 0 | 88.7 | 2 |
| VP72.7 | 3 | 100,895 | 13,306 | 20.0 | 4.3 | 0 | 85.9 | 3 |
| VP72.6 | 3 | 94,364 | 24,015 | 18.0 | 3.7 | 0 | 83.9 | 4 |
| VP72.5 | 3 | 115,693 | 19,044 | 32.0 | 19.2 | 0 | 66.1 | 5 |
| VP72.1 | 3 | 99,933 | 12,401 | 22.7 | 4.2 | 3 | 23.1 | 6 |
| VP72.3 | 1 | 0 | 0 | 0 | 0 | 0 | 0 | 7 |

Table S7. Cost of goods (COG) to make 1 ASFV test kit

| S.N. | Parts Name | Vendor | Units | Price/ Unit | Total Price (USD) | Notes |
| --- | --- | --- | --- | --- | --- | --- |
| 1 | [1.5 mm Acrylic for cartridge](https://www.amazon.com/Pieces-Acrylic-Sheet-Transparent-Project/dp/B09MRFHFTR/ref=sr_1_3?crid=3FYCOEH450GSN&dib=eyJ2IjoiMSJ9.ihSLCOsV10Aqr9Z5-Uye32nv9oSz0R-J5qUYbwjWWMQKYl1RgDehqXhzLbU7mufas7CLMOpU2WFHRfz94CUyVGi14zTMnIo5BeQKWOlkLHeVmG_6tsV7LUgD_7xzA6hRAMBOZljm_7sD73SNH3t86W8CtePxxN_u9X_Zn_P7ZMLPCiZ_OFJcvoyUShRyZRQDPykD06s-K0KOe1wCPLy_jfd7g_HYzriw5n3z3lNHrBs.Uh2vt7sTtHesFMpOyfvu4WRSIXqEJfbe9EZxcD4kRqk&dib_tag=se&keywords=1.5%2Bmm%2Bblack%2Bacrylic&qid=1756158730&sprefix=1.5%2Bmm%2Bblack%2Bacrylic%2Caps%2C111&sr=8-3&th=1) (Outus, ASIN: B09MRFHFTR) | Amazon | 1/200 | 4.3 | 0.02 | One 12-inch x 12-inch acrylic panel can be used for fabricating 200 acrylic cartridge |
| 2 | [Adhesive PCR Plate Seals](https://www.thermofisher.com/order/catalog/product/AB0558) (Catalog number: AB0558) | ThermoFisher Scientific | 1/1,000 | 188.65 | 0.19 | 1/10 full PCR tape is required (both sides) and it can be bought at a bulk of 100 units. |
| 3 | [LAMP assay for µPADs](https://www.sciencedirect.com/science/article/pii/S0956566325005640) | Multiple vendors | 2 | 0.74 | 1.48 | Cost estimate from (Ahmed et al., 2025b) |
| Total cost of 1 ASFV test kit | | | | | **1.69** |  |
