## Supplementary material for "A rapid, field-deployable paper-based biosensor for the detection of African swine fever virus in whole blood": ASF_Field_Manual.docx

**African Swine Fever Virus Diagnostics Test Manual 1.0**

This guide explains how to use a paper-based colorimetric DNA detection test developed in Dr. Mohit Verma’s lab at Purdue University in West Lafayette, Indiana. The test checks for synthetic *African Swine Fever* virus DNA present in blood. After the test runs, you'll be able to determine if the viral DNA is present in the sample.

Safety precautions: Please wear a pair of gloves provided with the kit. Always ensure proper safety and hygiene practices are followed when handling samples to prevent contamination.

**Components of the kit:**

Cartridge holder

Heating rod

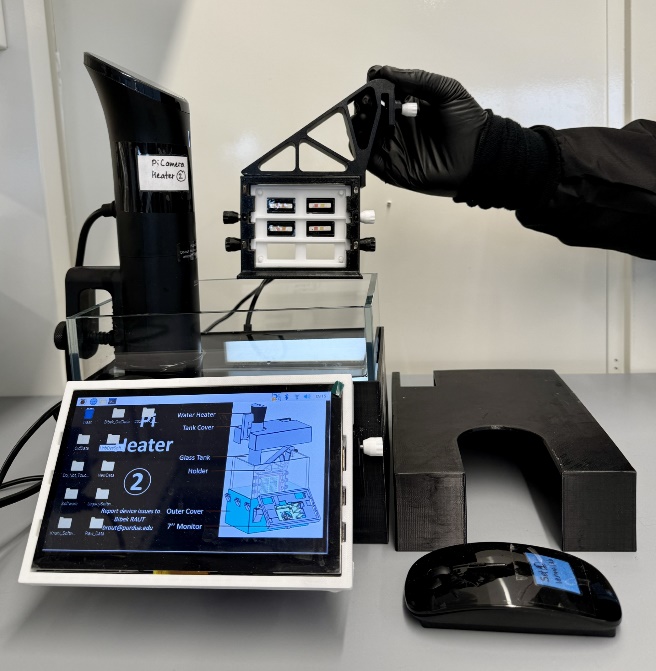

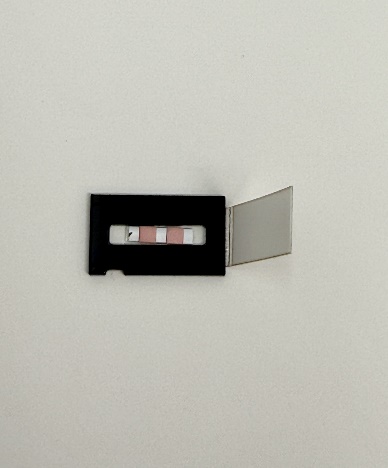

**Two paper pads in one ASFV diagnostic chip**

Sealing tape

Left pad is No Primer Control (NPC) and is indicated with a black dot near the pad

Right pad is with primer for detection

Monitor

Mouse

Water tank

Heater lid

Back Light

**Heater with cartridge holder**

**Cartridge chips containing the reaction pads to load the template with sealing tape attached**

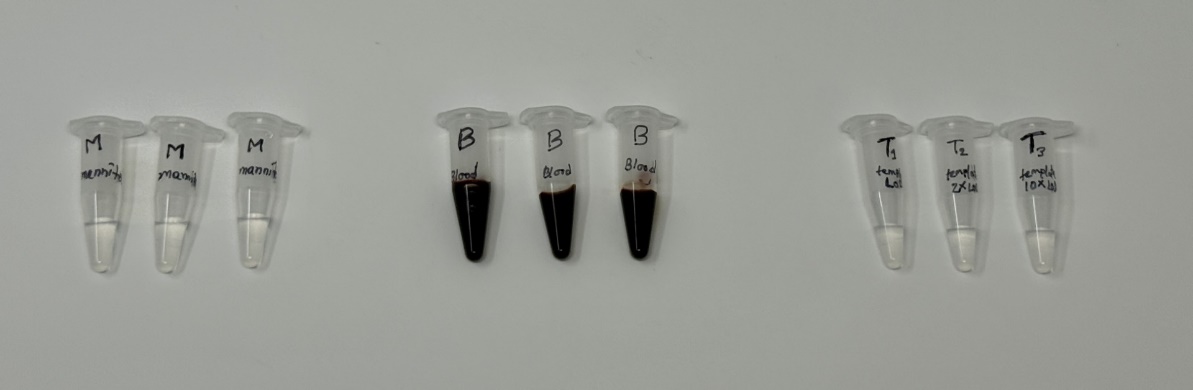

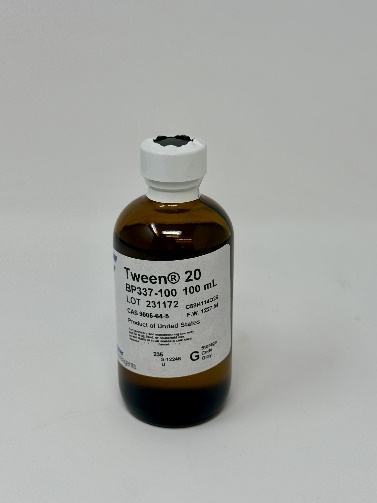

4 Tubes “B” (With 490µL whole blood in each)

3 Tubes - T1, T2, T3 (With synthetic DNA of ASF virus)

1 bottle of Tween 20

4 Tubes “M” (With 950 µL D-mannitol solution in each)

T2 for 2X LoD

T3 for 10X LoD

T1 for 1X LoD

**Reagents and Templates**

**Step-by-step guide**

1. **Heater preparation and software launch:**

- Fill the glass tank with water up to the point covered with the surrounded black cover.
- Add around 1 cap full of Tween 20 solution in the water.
- Attach the heater rod to the left side wall, then press the button on the heating rod to turn on the heating.
- The temperature is preset to 149.5°F (65°C). Let the temperature increase to the preset temperature as you prepare the devices for loading into the heater.
- Turn on the light placed at the back of the heater and keep it connected to the power during the experiment.
- On the monitor screen, browse to the Desktop/Software_ASF_Heater folder.
- Double-click PaperLAMP_ASF_final.py to open it. Keep it open to ensure there is no delay in starting the imaging right after loading the cartridge chips.

2. **Synthetic ASF Virus DNA Template Preparation:**

- There are 3 tubes with different concentrations of DNA in the package **(T1= 3.33×10^3^copies/µL, T2= 6.66×10^3^copies/µL and T3= 3.33×10^4^copies/µL)** comparable with 1X LoD (Limit of Detection), 2X LoD and 10X LoD in the final reaction (the limit of detection is 25copies/reaction).
- Make sure the templates are thawed properly before using (Thawing is done in room temperature)
- After the template is thawed properly using a pipette mix the template to ensure homogeneous mixture of the template.

3. **Spike blood with DNA:**

- Sample preparation for 1X LOD: Take 10 µL of template from tube “T1” and add that with the 490 µL of blood present in any of the tubes “B”. You can mark the tube as “B1”.
- Sample preparation for 2X LOD: Take 10 µL of template from tube “T2” and add that with the 490 µL of blood present in any other tube “B”. You can mark the tube as “B2”.
- Sample preparation for 10X LOD: Take 10 µL of template from tube “T3” and add that with the 490 µL of blood present in another tube “B”. You can mark the tube as “B3”.

4. **Blood Dilution with D-mannitol to get 5% blood:**

- After mixing the template with blood, take 50 µL of the spiked blood from tube “B1” in one of the tubes M that holds 950 µL of D-mannitol solution. Mix the blood with the mannitol solution properly. This will create the blood ready to be tested for 1X LoD. You can mark the tube as “M1”.
- Repeat the same step with the blood from tubes “B2” and “B3” to get the blood ready for 2X LoD and 10X LoD respectively. You can mark them as “M2” and “M3”.
- Now, you have the 5% blood ready in three different concentrations (i.e., tube “M1” for 1X LoD, “M2” for 2X LoD and “M3” for 10X LoD).
- After preparing the 3 concentrations, take 50 µL of blood **(which is not spiked with DNA)** from the unused tube “B” and mixed that with the last unused tube M. This will create the No Template Control (NTC) solution.

5. **Releasing the sample on the chip:**

- Pipetting the diluted blood: With the help of a pipette, take 7.5 µL of the diluted blood (see Figure 1).
- Release the sample: Release the blood sample on one reaction pad included in the chip. Repeat the same process for the other paper pad.
- Seal the chip: Peal the extended part of the tape and close the chip (see Figure 2). Make sure that the sealing is proper by pressing the edges.
- Start fresh: Use a new tip to load the template in each reaction pad in every new chip to prevent cross-contamination.

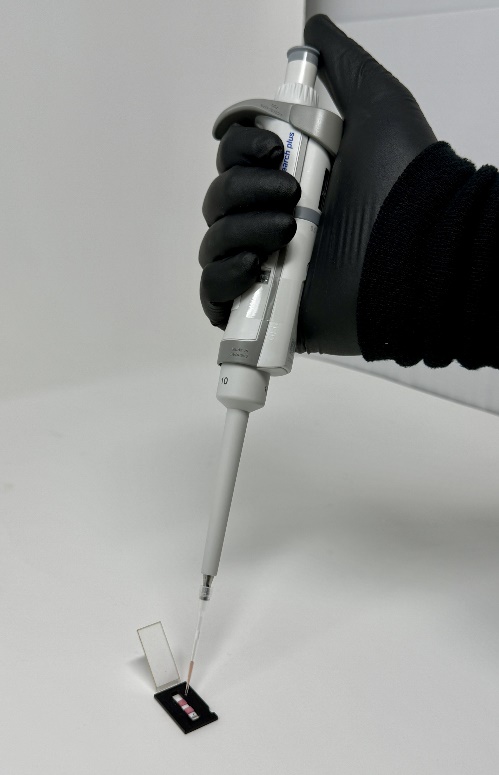

**Figure 1**

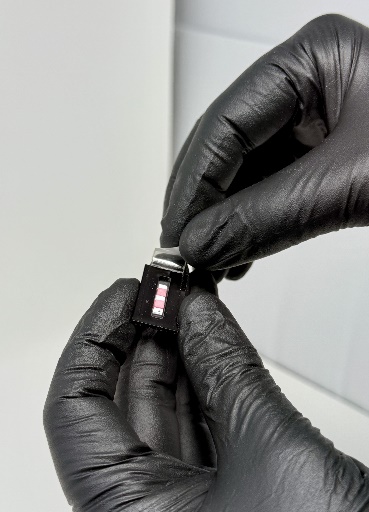

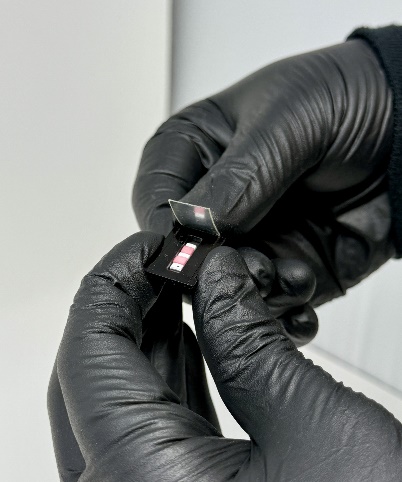

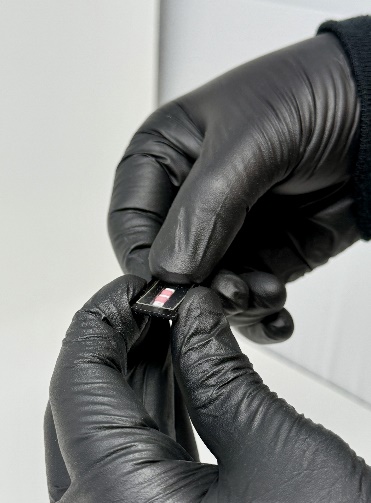

C

B

A

**Figure 2**

6. **Loading the chips on the frame:**

- Take the back part of the cartridge frame (see Figure 3A).
- Place the cartridge chip over the designated spot on the frame. Each frame can hold up to six chips (see Figure 3B).
- After putting the chips (up to six chips) take the front part of the frame and put that over the back part of the frame, on top of the chips (see Figure 3C). Make sure to keep note of which chip is placed where in the frame for future reference.

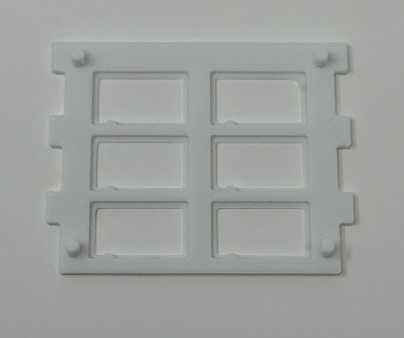

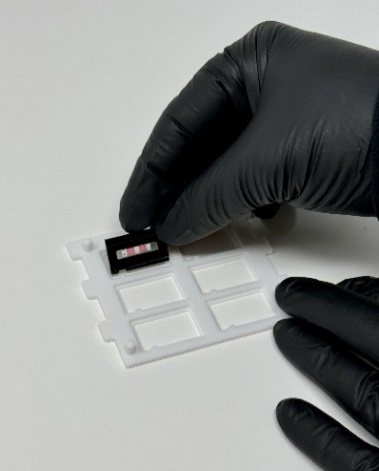

B

A

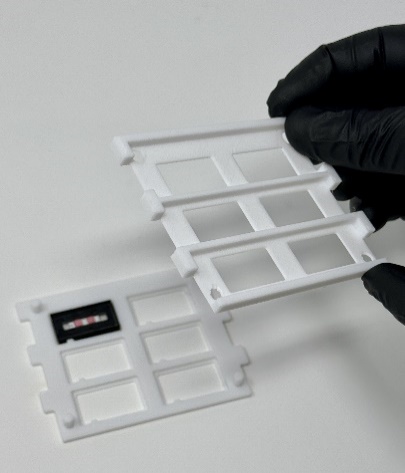

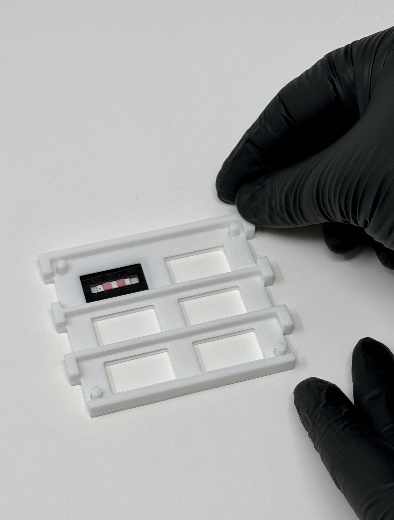

C

D

**Figure 3**

7. **Setting up the cartridge holder:**

- Position the cartridge frame over the cartridge holder as demonstrated in Figure 4A.
- Put the cartridge frame inside the cartridge holder (Figure 4B).
- Lock the position of the cartridge frame by securely tightening the knobs (Figure 4C). The screw should go over the frame to secure the frame to the holder.

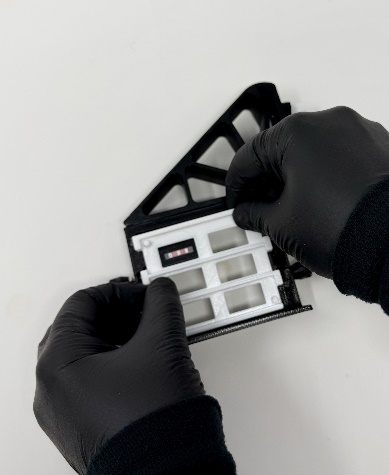

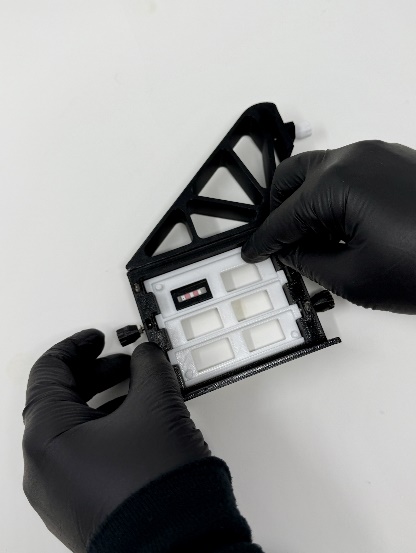

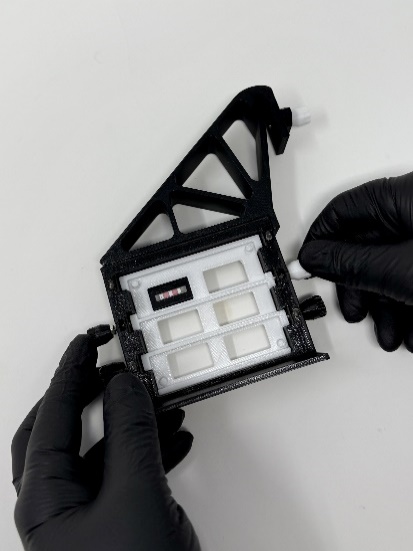

C

B

Anti-clockwise

Clockwise

Putting the frame inside holder

A

**Figure 4**

8. **Placing the holder in the heater:**

- Insert the holder with the cartridge into the heater as shown in the picture provided (Figure 5A).
- Secure the cartridge holder by slightly tightening the knob to the heater (Figure 5B). Do not tighten too much else it will break the holder. Tighten enough to secure the holder.
- Put on the heater lid.

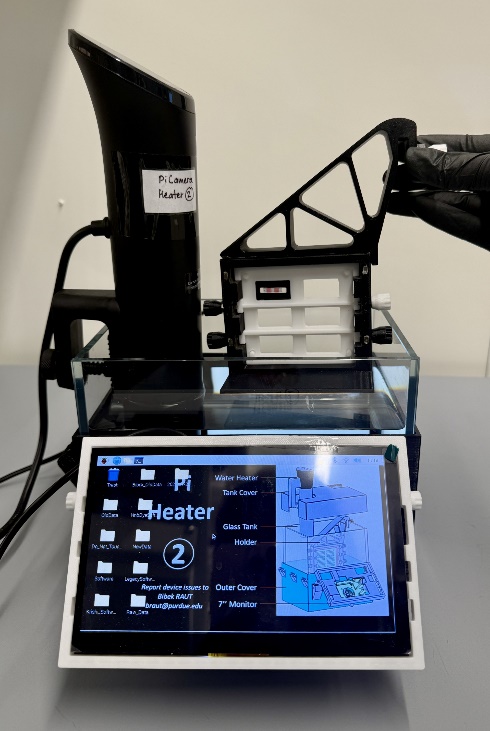

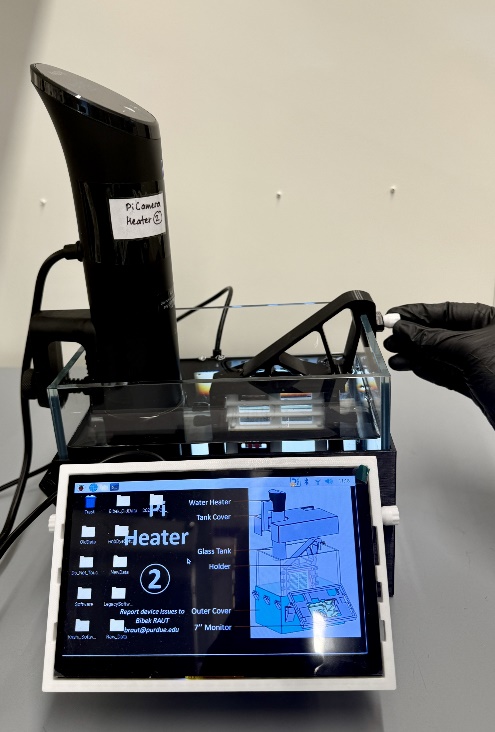

A

B

Clockwise

**Figure 5**

9. **Starting the software:**

- Press the Play Icon
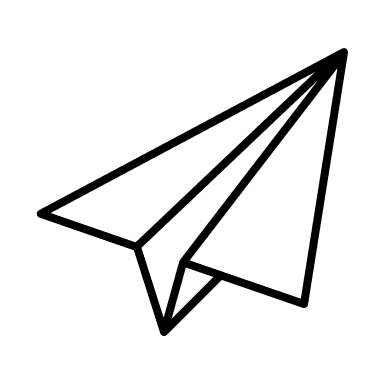
 icon to start data capture.
- Wait for 60 minutes for the reaction to complete.

10. **Getting the results**

- Click “Summary Report” to view the results.
- Press “Stop” to end the session.

Notes: As a resolution of any unexpected mistake, the entire kit has extra 10 cartridge chips, 1 tube of D-mannitol solution and 1 tube of whole blood. It is recommended to store the templates at -20°C and Blood and D-mannitol at 4°C.
